# Signaling mechanisms and agricultural applications of (*Z*)-3-Hexenyl Butyrate-mediated stomatal closure

**DOI:** 10.1101/2023.05.26.542418

**Authors:** Celia Payá, Borja Belda-Palazón, Francisco Vera-Sirera, Julia Pérez-Pérez, Lucía Jordá, Ismael Rodrigo, José M Bellés, M Pilar López-Gresa, Purificación Lisón

## Abstract

Biotic and abiotic stresses can severely limit crop productivity. In response to drought, plants close stomata to prevent water loss. Besides, stomata are considered the main entrance of several pathogens. Therefore, the development of natural products to control stomata closure can be considered a sustainable strategy to cope with stresses in agriculture. Plants respond to different stresses by releasing volatile organic compounds (VOCs). Green leaf volatiles (GLVs), which are commonly produced across different plant species after tissue damage, comprise an important group within VOCs. Among them, (*Z*)-3-hexenyl butyrate (HB) was described as a natural inducer of stomatal closure, playing an important role in stomatal immunity, although its mechanism of action is still unknown. Here, through different genetic, pharmacological, and biochemical approaches, we uncover that HB perception initiates various defense signaling events such as activation of Ca^2+^ permeable channels, mitogen-activated protein kinases (MPKs) and production of NADPH oxidase-mediated reactive oxygen species (ROS). Furthermore, HB-mediated stomata closure resulted to be independent of abscisic acid (ABA) biosynthesis and signaling. Additionally, exogenous treatments with HB alleviate water stress and improve fruit productivity in tomato plants. The efficacy of HB was also tested under open field conditions, leading to enhanced resistance against *Phytophthora* spp. and *Pseudomonas syringae* infection in potato and tomato plants, respectively. Taken together, our results provide insights into HB signaling transduction pathway, confirming its role in stomatal closure and plant immune system activation, and proposing HB as a new phytoprotectant for the sustainable control of biotic and abiotic stresses in agriculture.

## Introduction

Plants have evolved a complex and efficient innate immune system to cope with pathogen attacks. Two different defensive layers are involved in plant immune responses ^1^. The first line of plant innate immunity is established upon the recognition by receptors on the plasma membrane (pathogen recognition receptors; PRRs) of conserved microbial features, called as pathogen- and microbe-associated molecular patterns (PAMPs and MAMPs), activating the PAMP-triggered immunity (PTI) ^2,3^. At this level, through different PRRs, plants are also able to perceive molecules known as damage molecular patterns (DAMPs), released from plant cells upon pathogen attack. This specific recognition also triggers a PTI response known as DAMP-triggered immunity (DTI) ^4^. The second layer is started by the recognition of virulence factors, also called effectors, by intracellular receptors (nucleotide-binding, leucine-rich repeat receptors; NLRs), activating the effector-triggered immunity (ETI) ^5,6^. Although both types of plant immune responses involve different activation mechanisms, PTI and ETI share many downstream components and responses, such as reactive oxygen species (ROS) production, increases in cytosolic calcium (Ca^2+^) levels or activation of mitogen-activated protein kinase (MPK) cascades. The amplitudes and dynamics differ in each defensive level, postulating that ETI is an “accelerated and amplified PTI” ^7^.

Many foliar pathogens, including bacteria, are unable to directly penetrate plant tissues and they use natural openings such as stomata. As a countermeasure, plants have evolved a mechanism to rapidly close their stomata upon PAMPs/MAMPs perception limiting bacterial entrance, a defensive response known as stomatal immunity ^8^. The bacterial MAMP flg22, an immunogenic epitope of the bacterial flagellin, is recognized by the PRRs receptor FLAGELLIN-SENSITIVE2 (FLS2) and the coreceptors BRI1-ASSOCIATED RECEPTOR KINASE1 (BAK1) and BOTRYTIS-INDUCED KINASE 1 (BIK1) ^9^. This recognition by the FLS2 complex triggers a cascade of signaling events including increases of Ca^2+^ influx, ROS burst through the activation of plant NADPH oxidases, MAPK cascades and activation/inhibition of ion channels, all leading to stomatal immunity ^10^.

The phytohormone abscisic acid (ABA) is essential in stomatal defense during plant immunity ^11^. In fact, ABA-induced stomatal closure shares a common signaling pathway with PAMP-induced stomatal closure, including ROS burst, nitric oxide (NO) intermediate accumulation, activation of S-type anion channels, or the inhibition of K^+^ channels ^10^. Despite this, there are some differences in the canonical stomatal immunity signaling components in both PTI and ABA signaling pathways. ABA-induced ROS production depends on the kinase OST1, which phosphorylates and activates the NADPH oxidases RBOHF/RBOHD. However, during PTI stomatal closure, ROS are produced by the activation of RBOHD that is directly phosphorylated by the plasma-membrane-associated kinase BIK1 ^9,10^. Furthermore, MPKs are known to play a specific role in stomatal immunity. MPK3 and MPK6 positively regulate flg22-triggered stomatal closure in a partially redundant manner, but they are not involved in ABA-mediated stomatal closure ^12,13^.

Volatile organic compounds (VOCs) act as fast signaling molecules that activate plant defensive response pathways between distant organs, and they even allow communication between plants ^14–16^. Several studies have demonstrated that VOCs are able to induce defenses against herbivorous insects, pathogens, and even environmental stresses ^15,17–19^. Defense priming against pathogens induced by VOCs has been considered as a sort of “green vaccination” ^20^.

A non-targeted GC-MS metabolomics analysis revealed the VOCs profile associated with the immune response of tomato cv. Rio Grande plants upon infection with either virulent or avirulent strains of the model bacterial pathogen *Pseudomonas syringae* pv. tomato (*Pst*) DC3000. In the case of the avirulent infection which leads to the establishment of ETI, the VOC profile of immunized plants was characterized by esters of (*Z*)-3-hexenol with acetic, propionic, isobutyric or butyric acids, and several monoterpenoids such as linalool or α-terpineol ^21^. The defensive role of these compounds was tested trough exogenous application in tomato plants and, among all these compounds, treatments with (*Z*)-hexenyl butyrate (HB) resulted in the transcriptional upregulation of defensive genes, stomatal closure and enhanced resistance to the bacterial infection with *Pst* DC3000, confirming the role of HB as a natural defense elicitor ^22^. Additionally, HB-mediated stomatal closure was effective in different plant species, which also led to accelerated ripening in *Vitis vinifera* ^22,23^.

In this study, we investigated the signaling mechanisms underlying HB-mediated stomatal closure and plant immunity. Furthermore, we tested the efficacy of this compound in water stress and its effectiveness was also assessed against both biotic and abiotic stresses under field conditions. Considering all our results together, we propose a new role for HB as a natural elicitor against biotic and abiotic stresses.

## Results

### Treatments with HB induce transcriptomic changes related to plant immunity and photosynthesis

To provide insight into the molecular mechanisms underlying the tomato plant response to HB, an RNA-seq analysis was performed on leaf tissues from mock and HB-treated tomato plants. After trimming and filtering data, a total of 1456 genes were found to be differentially expressed (DEGs) between HB- and control-treated plants (DEG threshold = 2), indicating that HB treatments produce an outstanding effect at the transcriptional level. To classify DEGs, a GO enrichment analysis for biological processes (BP), cellular components (CC) and molecular functions (MF) was performed for up- and down-regulated DEGs, independently.

Among the upregulated genes, 1122 DEGs were annotated, being the BP category the most enriched one (Figure 1A). In this regard, most of the DEGs were putatively involved in organonitrogen compound metabolic processes (GO:1901564), chitin metabolic process (GO:0006030) or amino glycan and amino sugar metabolic processes (GO: 0006040, GO:0006022). MF category included chitinase activity (GO:0004568), chitin binding (GO:00080761) or endopeptidase activity (GO: 0004175), which are mainly related to pathogen defensive responses. On the other hand, only 334 downregulated DEGs were annotated, and most of them were associated with the CC category mainly related to plastoglobule (GO:0010287), photosystem (GO:0009521) and chloroplast (GO:0009507) (Figure 1B). Indeed, the most enriched groups when biological processes were assessed were related to photosynthesis (GO:0015979) and the only enriched MF term was chlorophyll binding (GO:0016168).

**Figure 1.**
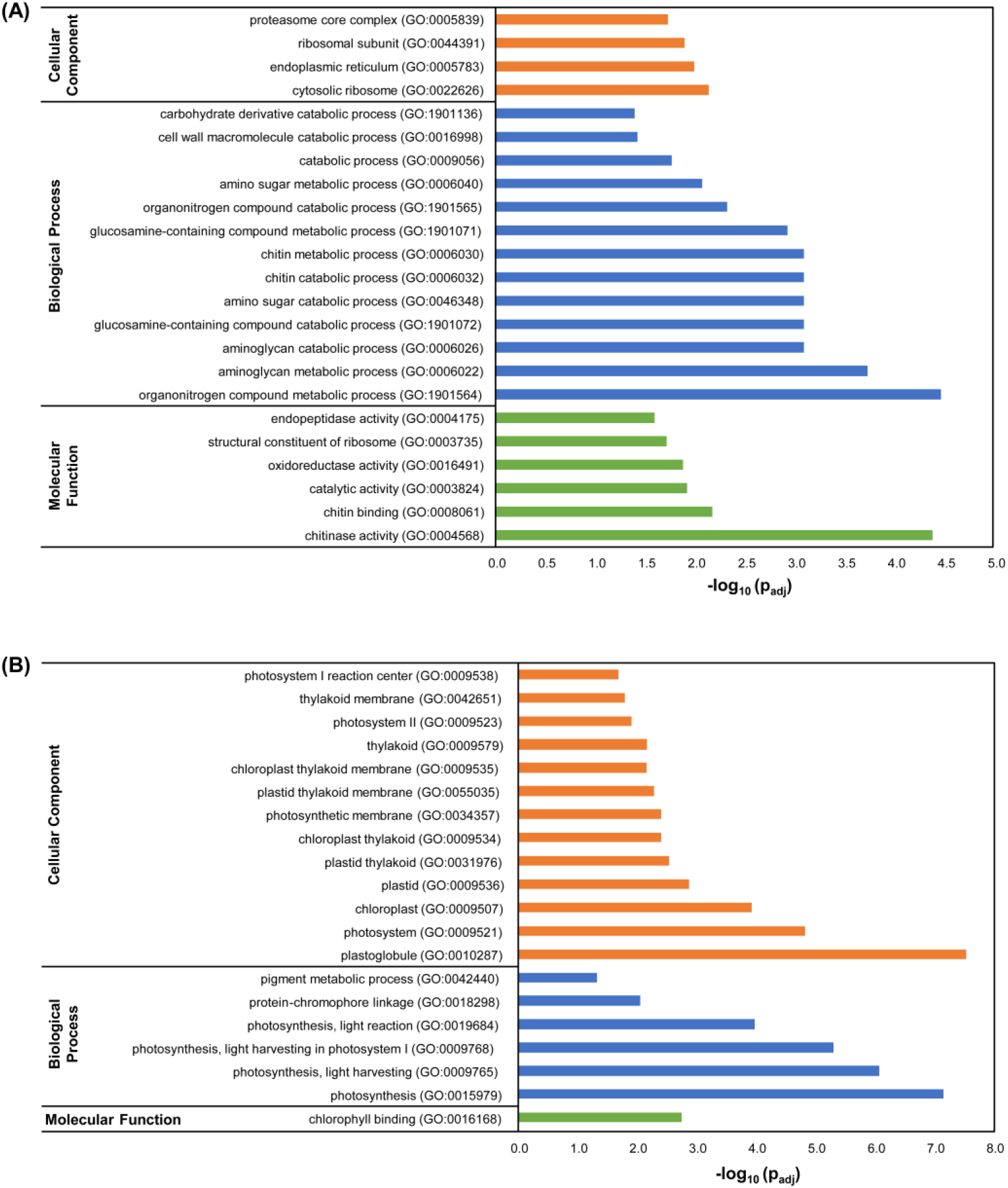
Transcriptomic analysis suggests a role of HB in plant immunity. Functional profiling analysis of upregulated (A) and downregulated (B) DEGs between HB- and control-treated plants.

Both defensive and photosynthesis-related gene expression were confirmed by RT-qPCR (Supplemental Figure 1), observing a statistically significant induction for the pathogenesis-related genes *PR1* (X68738) and *PR5* (X70787), as well as a gene coding for a putative PRR immune leucine-rich repeat receptor-like kinase (Solyc12g036793), and also a gene involved in heavy metal resistance and detoxification (Solyc06g066590). Besides, a reduction in the expression of genes encoding chlorophyll binding proteins (Solyc06g069730; Solyc06g069000; Solyc12g011280) was also validated. Therefore, RNA-seq transcriptomic analysis revealed that HB treatments induced plant defensive responses against pathogens as well as photosynthesis reprogramming.

### ROS production via NADPH oxidases and Ca^2+^ signaling, but not ABA, are required for HB-mediated stomatal closure

To decipher the signaling pathway by means of which HB mediates stomatal closure, a suitable method was developed for stomatal measurement analyses (Materials and Methods). In these experiments, we studied the effect of HB in comparison with flg22 and ABA, the positive controls for stomatal closure, considering both dose-response and time-course assays of these compounds to optimize the different treatments (Supplemental Figure 2). Once set up, we studied whether ABA was necessary for the HB-dependent stomata closure. For this purpose, we took advantage of both *flacca* (*flc*) tomato mutants that are impaired in ABA biosynthesis and their corresponding parental Lukullus wildtype (WT) plants ^24^. As expected, HB, flg22 and ABA treatments promoted stomatal closure in WT plants. In *flc* mutants, the stomatal aperture ratio in mock conditions was significantly higher compared to WT plants, probably due to a reduction of the ABA levels. However, ABA treatments in *flacca* mutants closed the stomata to the same extent than flg22 and HB, suggesting that ABA biosynthesis is not required for flg22- and HB-mediated stomatal closure (Figure 2A).

**Figure 2.**
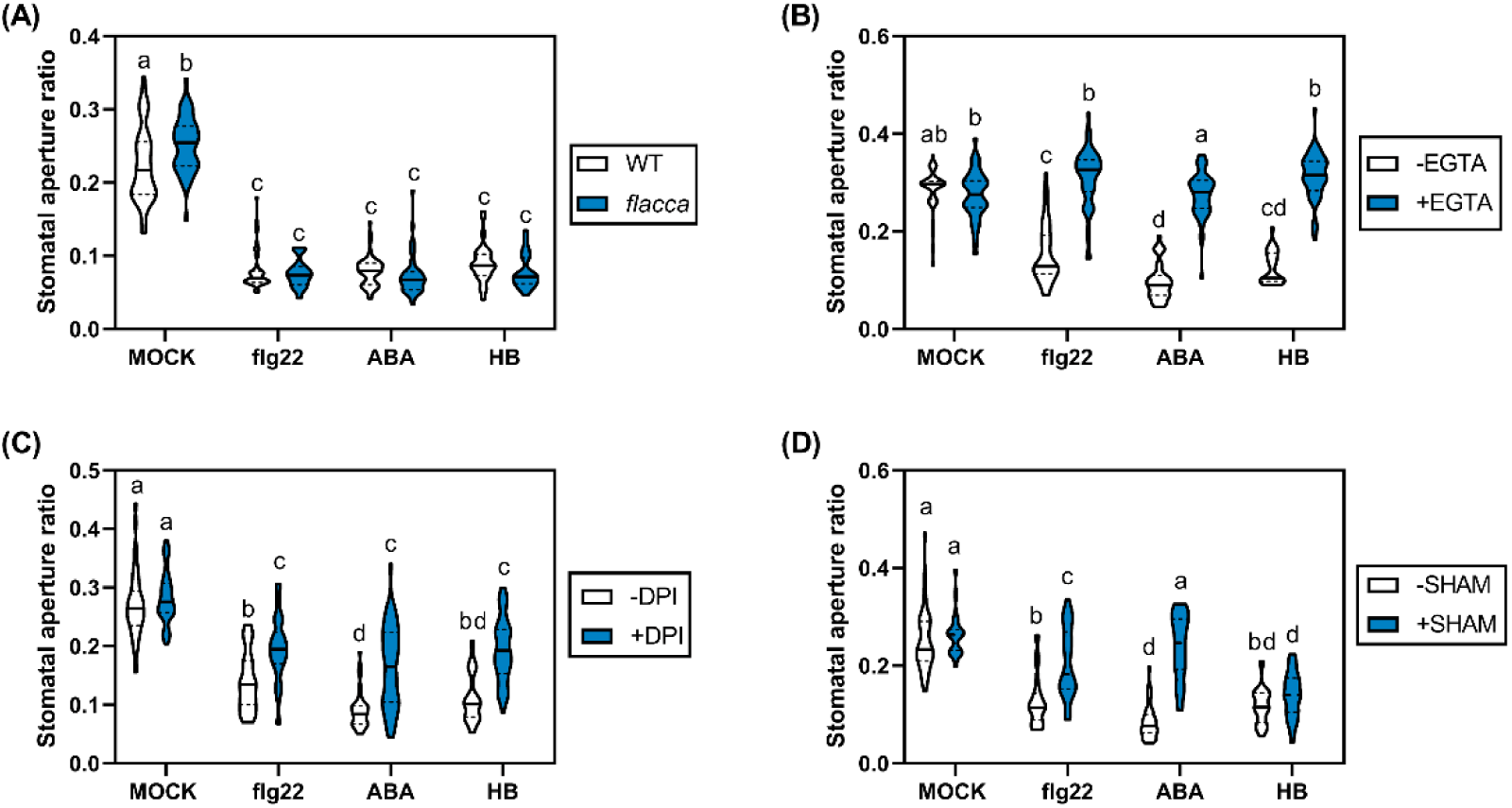
HB-mediated stomatal closure requires Ca2+ signaling and ROS production by NADPH oxidases, but not ABA biosynthesis. (A) Lukullus (WT) and *flacca* tomato leaf discs were floated on liquid MS for 3 h under light. Then, 1 μM flg22, 10 μM ABA or 50 μM HB were applied, and stomatal aperture ratio was determined 2 h after treatments. In the case of chemical inhibitors experiments, 2 mM EGTA (B), 20 μM DPI (C) or 2 mM SHAM (D) were added in the liquid medium for MoneyMaker tomato leaf discs before elicitor treatments. Violin plots represent the stomatal aperture ratio of 50 stomata for each treatment. Different letters indicate statistically significant differences for each genotype and treatment (*p* < 0.05, two-way ANOVA with Tukey HSDTest).

To analyze the role of Ca^2+^ and ROS in the HB-mediated stomatal closure, we performed pre-treatments with the specific Ca^+2^ ion chelator EGTA, and the ROS production inhibitors DPI (Diphenyleneiodonium, which inhibits NADPH oxidase) or SHAM (salicylhydroxamic acid, acting on peroxidases) before triggering the stomatal closure with either HB, flg22 or ABA. Thereby, EGTA pretreatments completely abolished the stomatal closure mediated by all HB, flg22 and ABA treatments, indicating that Ca^2+^ signaling is essential for stomatal closure (Figure 2B). Interestingly, pretreatments with DPI partially abrogated the stomatal closure induced by HB, flg22 and ABA, suggesting that NADPH-dependent ROS production is somewhat necessary for this process (Figure 2C). However, unlike flg22 and similar to ABA treatments, SHAM did not inhibit HB-induced stomatal closure, indicating that stomatal closure promoted by HB is independent of ROS production by peroxidases (Figure 2D).

Our results appear to indicate that HB-mediated stomatal closure requires Ca^2+^ fluxes and is partially dependent on ROS generation by NADPH oxidases but is nevertheless independent of ABA biosynthesis and ROS generated through peroxidases.

### Activation of MPK3 and MPK6 is essential for HB-mediated stomatal immunity

To test whether HB intracellular signaling occurs through activation of MPK-cascades, we analyzed the phosphorylation of MPK3 and MPK6 in tomato and *Arabidopsis thaliana* leaf discs after treatments with flg22, ABA and HB. Regarding tomato samples, HB induced MPK3 phosphorylation 15 minutes after treatments, and this activation persisted until 60 minutes after treatments (Figure 3A). Nevertheless, in *Arabidopsis thaliana* tissues, MPK3/6 phosphorylation was observed at 60 minutes after HB treatments, indicating that MPKs activation phenomenon occurs earlier in tomato than in *Arabidopsis thaliana* plants (Figure 3B).

**Figure 3.**
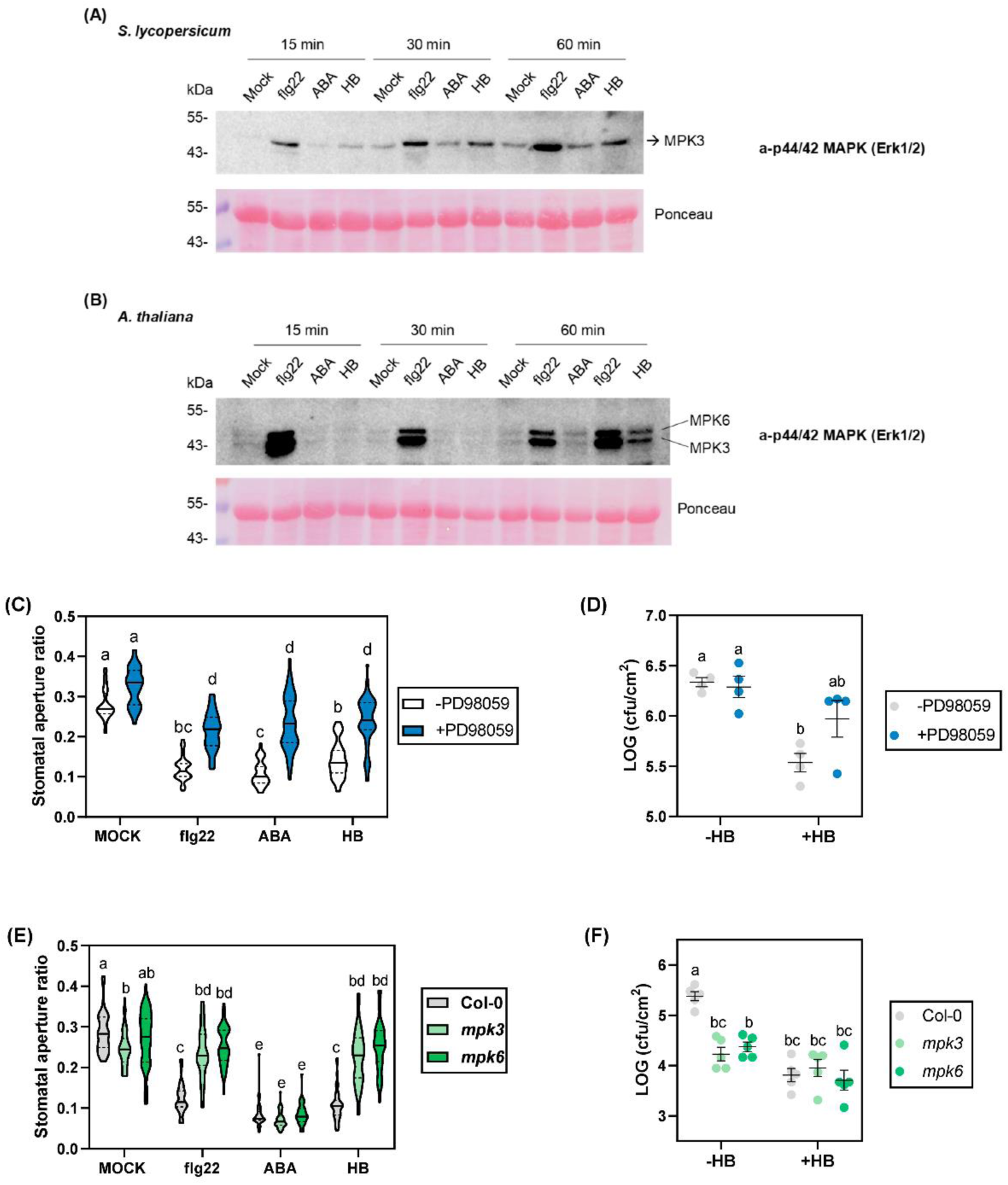
Activation of MPK3 and MPK6 is involved in HB-mediated stomatal immunity. MPK activation assay in tomato (A) and *Arabidopsis thaliana* (B) leaf discs 15, 30 and 60 minutes after treatments with 1 μM flg22, 10 μM ABA and 50 μM HB. MPK activation was detected by immunoblot analysis using the Phospho-p44/42 MAPK (Erk1/2; Thr-202/Tyr204) rabbit monoclonal antibody. Western blot experiments were performed three times and yielded similar results. (C) Stomatal aperture ratio was measured in tomato leaf discs floated on MS liquid for 3 h under light, pre-treated with the MPKs inhibitor PD98059 20 μM and followed by treatments with 1 μM flg22, 10 μM ABA and 50 μM HB for 2 h. (D) Growth of Pst on tomato leaves of control (-HB) and HB-treated (+HB) after treatments with the MPKs inhibitor PD98059. Plants were sprayed with with PD98059 100 μM for 3 hours, subsequently treated with HB 5 μM or water for 24 h into methacrylate chambers, and then dip inoculated with Pst. Bacterial growth measurements were done 24 h after inoculation. (E) Stomatal aperture ratio of Arabidopsis Col-0, mpk3 and mpk6 mutants leaf discs 2 h after treatments with flg22, ABA and HB. (F) Growth of Pst on Arabidopsis thaliana leaves of control (-HB) and HB-treated (+HB) 3 days after inoculation. Plants were treated with HB 5 μM or water for 24 h into methacrylate chambers, and then infected by spray with Pst. In stomatal aperture experiments, 50 stomata were measured per each treatment and condition (violin plots). Data correspond to at least four independent plants ± SEM of a representative experiment. Statistically significant differences are represented by different letters (*p* < 0.05, two-way ANOVA with Tukey HSD).

To confirm the importance of MPK3/6 activation in HB-mediated stomatal immunity, pre-treatment with the MPKs inhibitor PD98059 on tomato leaf discs was performed. After PD98059 application, flg22, ABA and HB treatments diminished their ability to induce stomatal closure, confirming that MAPKs play a pivotal role in all the elicited stomatal responses (Figure 3C). Furthermore, treatments with PD98059 were also carried out *in planta* (see Materials and Methods). In this regard, PD98059 application diminished the HB enhanced resistance to bacterial infection (Figure 3D). These results indicate that HB induces stomatal immunity partially via activation of the MPK3/6 signaling cascades in tomato plants.

To better characterize this, a genetic approach was performed in *Arabidopsis thaliana mpk3* and *mpk6* mutants analyzing the stomatal behavior as described in Materials and Methods. As expected, ABA treatments dramatically reduced stomatal aperture in both Arabidopsis mutants and WT plants. In contrast, and in accordance with the chemical inhibitor experiments performed in tomato plants, stomata aperture ratio was identical to that observed in mock-treated plants not only in flg22-but also in HB-treated *mpk3* and *mpk6* mutants, whilst treatments were efficient in the corresponding WT Arabidopsis plants (Figure 3E). The incapacity of HB in closing stomata in *mpk3* and *mpk6* mutants resulted in non-enhanced resistance to bacterial infection with *Pst* after HB treatments (Figure 3F).

Additionally, the expression of MPK3/6 target genes was also explored upon HB treatment. As shown in Supplemental Figures 3A and 3B, HB treatments of tomato plants significantly induced *SlWRKY33A* and *SlWRKY33B* transcription factors, which correspond to tomato orthologs of *AtWRKY33.* This gene has been described to work downstream of MPK3/MPK6 in the reprogramming of the expression of genes involved in camalexin biosynthesis and pipecolic acid in Arabidopsis ^25,26^. Since MAPK3/6 play a central role in the regulation of the ethylene response pathway ^27^, ethylene signaling-related genes like the ethylene receptor *ETR4* and an ethylene responsive factor (*ERF*) were also analyzed, observing a significant induction of the expression of both genes (Supplemental Figures 3C and 3D). The expression of these four genes was checked in the RNA-seq analysis (Supplemental Figure 3E), confirming the HB-mediated induction of the MPK3/6 target genes. Taken together, these results indicate that MPK3/6 are involved in HB-mediated stomatal immunity.

### HB treatments confer drought tolerance in tomato

The robust effect of HB in promoting the closure of stomata prompted us to study the effect of HB on drought tolerance. To that purpose, well-watered as well as drought-stressed tomato plants were periodically treated with HB, and stomatal opening and closing dynamics were monitored. Three days after drought exposure (DAD), stomatal aperture ratio in HB-treated plants was lower than in control-treated plants, confirming the role of HB as stomatal closure inducer in tomato plants, even in drought conditions (Figure 4A). It should be noted that HB treatments in well-watered conditions caused similar stomatal closure to those control-treated plants subjected to drought. Moreover, at 6 DAD, the stomatal behavior in all treatments and conditions was similar to that observed at 3 DAD, except in the control plants exposed to drought, in which it was not possible to take leaf samples due to the wilting phenotype of the plants.

**Figure 4.**
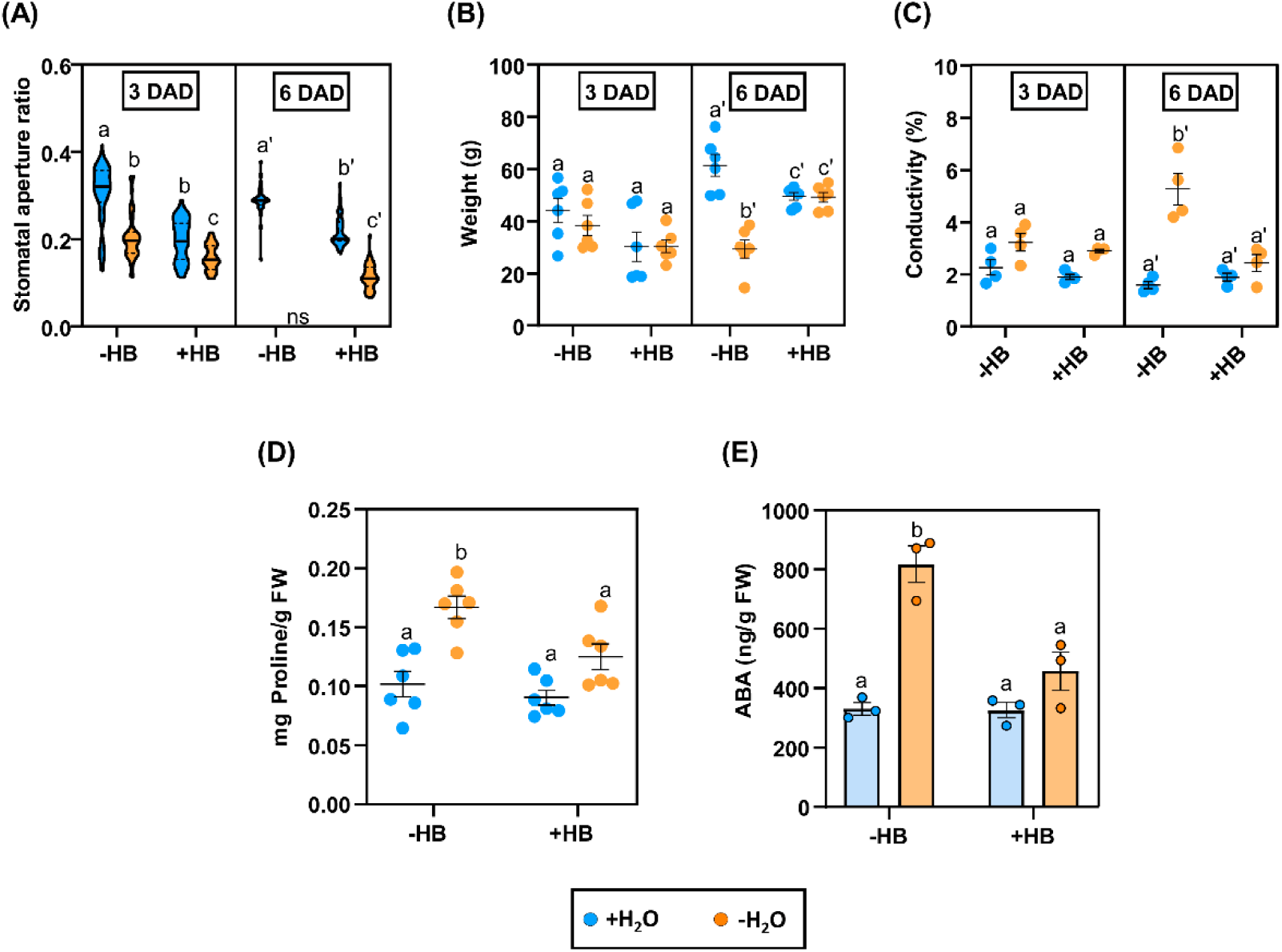
HB treatments induce tolerance to drought in tomato plants and reduce proline and ABA contents. Differences related to stomatal aperture ratio (A), weight (B), ion leakage (C), proline content (D) and ABA content (E) in tomato plants treated (+HB) or not (-HB) with HB, in normal (+H20; blue) or water stressed conditions (-H20; orange). Samples were taken 3 or 6 days after drought exposure (DAD). Data in (A) corresponds to 50 stomata per each treatment and condition (violin plots). Data in (B), (C) and (D) correspond to six independent plants ± SEM of a representative experiment. Data in (E) correspond to the averages of three independent plants ± SEM of a representative experiment. Different letters indicate statistically significant differences for each genotype and treatment (*p* < 0.05, two-way ANOVA with Tukey HSD).

To correlate the HB-mediated stomatal closure upon drought conditions with the water content of tomato plants, we also measured the fresh weight at 3 and 6 DAD. At 6 DAD, control-treated plants under drought conditions showed an approximately 50% reduction in the fresh weight. Nevertheless, in the case of HB-treated plants, no statistically significant differences were observed between well-watered plants and those that were under water deficit conditions (Figure 4B), thus indicating HB confers an outstanding drought tolerance.

Finally, plasma membrane damage was also evaluated by measuring electrolyte leakage in tomato leaves. In severe drought conditions (6 DAD), no significant differences were observed between HB-treated plants. However, control-treated plants subjected to drought displayed a higher percentage of ion leakage, suggesting the role of HB in cell membrane protection and stabilization (Figure 4C). Nonetheless, no differences in chlorophyll content were found between treatments and conditions, indicating that water-stressed plants had a withered appearance, but they were not completely collapsed (Supplemental Figure 4).

### Tomato plants treated with HB show lower proline and ABA levels under drought conditions

To better understand the role of HB in drought tolerance, levels of both proline, a well-known osmoprotectant, and ABA were analyzed in HB-treated or non-treated, and well-watered or non-watered tomato plants.

As expected, non-treated tomato plants subjected to water deprivation accumulated higher proline levels than well-watered plants. However, when HB-treated plants were analyzed, no statistical differences were found between non-stressed and water-stressed plants (Figure 4D). Accordingly, a similar trend was observed regarding proline biosynthesis, in which Δ^1^-Pyrroline-5-Carboxylate Synthetase 1 (P5CS1) is the rate-limiting enzyme ^28^. We observed that tomato *P5CS1* gene (*Solyc06g01970*) expression was significantly induced under drought conditions in both HB and non-treated plants. Nevertheless, *P5CS1* expression was lower in the case of HB-treated plants and the expression level was statistically similar to watered, non-treated plants (Supplemental Figure 5A).

Regarding the levels of ABA, no significant differences between untreated and HB-treated plants under well-watered conditions were observed. Remarkably, water-stressed plants treated with HB accumulated significantly less ABA than the stressed untreated plants, with levels similar to those observed in non-stressed plants (Figure 4E). The pattern of expression of genes that were involved in different critical steps of ABA biosynthesis, like 9-cis-epoxycarotenoid dioxygenase (NCED; Solyc07g056570), zeaxanthin epoxidase (Solyc02g090890), and ABA 8’-hydroxylase (Solyc04g078900) was also analyzed by RT-qPCR, observing a tendency to down-regulation upon HB treatments in irrigated plants (Supplemental Figure 5B, 5C and 5D). Besides, we analyzed the expression of marker genes of ABA signaling ^29^, such as Responsive to ABA 18 (*RAB18*), the transcription factor *MYB44,* and a late embryogenesis abundant (*LEA*) protein in tomato plants at 6 DAD (Supplemental Figure 6). Drought treatment induced the expression levels of *RAB18* and *LEA* in untreated and HB-treated plants. Interestingly, *LEA* gene induction in HB-treated plants appeared to be higher than in non-treated plants for both watered and drought stress conditions. The induction of *LEA* in water-stressed plants treated with HB (Supplemental Figure 6C), which accumulate lower levels of ABA (Figure 4E), appears to reinforce that HB induces the abiotic response in an ABA-independent manner.

Therefore, the effect of exogenous HB appears to alleviate drought stress, since the mechanisms that counteract water deprivation, including ABA and proline accumulation, are less activated in HB-treated plants.

### Transcriptomic changes in HB treatments under water stress revealed a higher induction of the abiotic responses and a repression of the water transport

To provide insight into the molecular responses of the plants exposed to HB subjected to water deprivation, a further RNA-seq analysis was performed. In a similar manner to the previous one, a GO enrichment analysis was performed for up- and down-regulated DEGs under water stress.

Among the upregulated genes, most of the DEGs were putatively involved in response to temperature stimulus and heat (GO:0009266 and GO:0009408), response to osmotic and salt stress (GO:0006970 and 0009651), response to abiotic stimulus (GO:0009628) or response to hydrogen peroxide and reactive oxygen species (GO:00042542 and 000302), which are mainly related to abiotic stress responses (Figure 5A). On the other hand, the annotated downregulated DEGs were associated with the fluid and water transport (GO:0042044 and 0006833) or with the response to hormone (GO:0009725) (Figure 5B).

**Figure 5.**
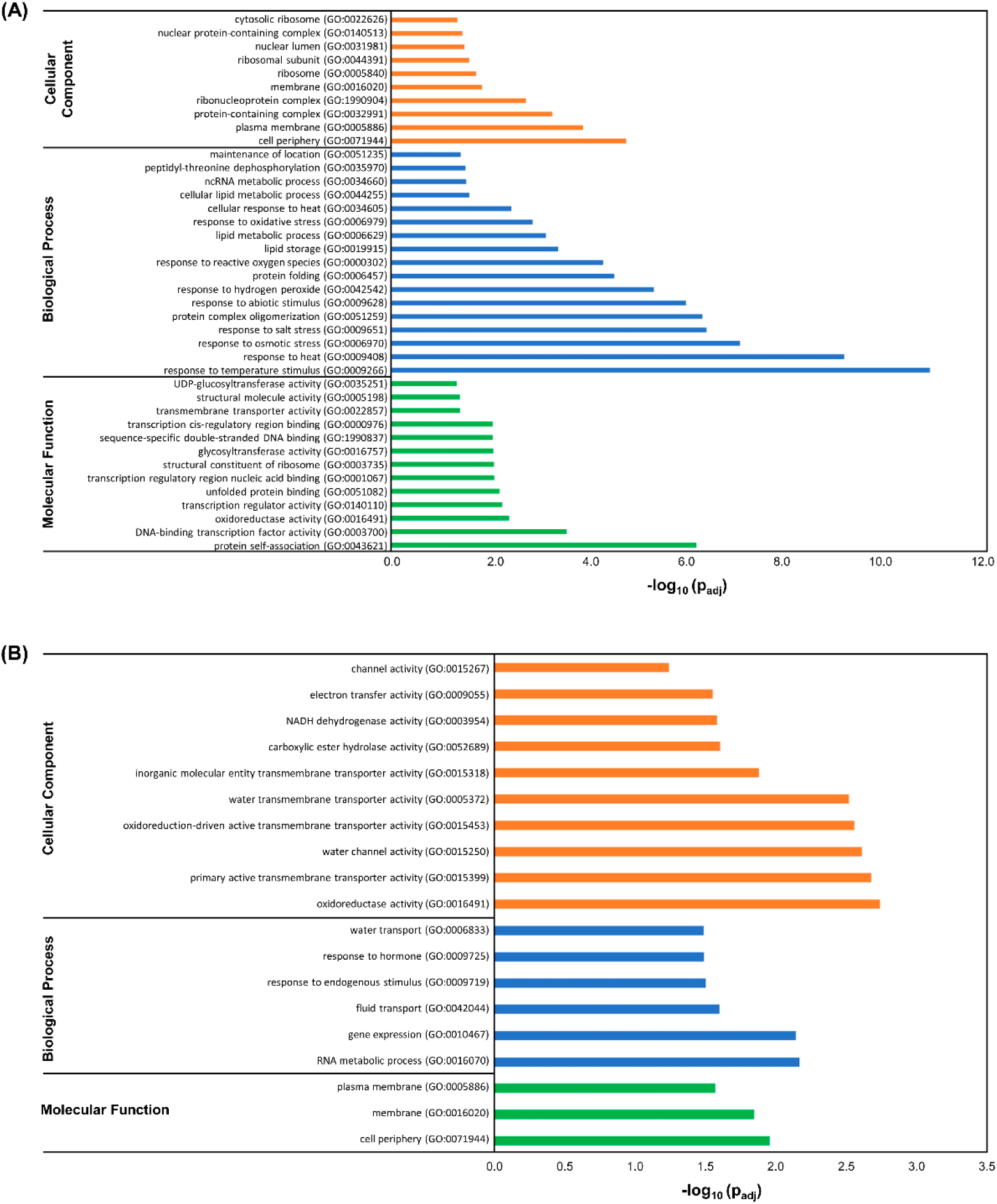
Transcriptomic analysis of HB-treated tomato plants under drought stress. Functional profiling analysis of upregulated (A) and downregulated (B) DEGs between HB- and control-treated plants under water deprivation.

Moreover, a gene clustering analysis with DEGS from both RNA-seq (DEG threshold = 2), was performed to discover groups of correlated genes between different treatments (non-treated and HB-treated) and conditions (well-watered and non-watered plants). Following this approach, 16 clusters were generated (Supplemental Figure 7). According to cluster trends, we grouped them in 3 groups of interest: first DEGS that share the same response to HB in well-watered and non-watered conditions (clusters 7, 10, 12 and 16), second DEGS that only respond to HB in non-abiotic stress conditions (clusters 5 and 6) and third DEGS that only respond to HB in water deficit conditions (clusters 2 and 3). When a GO analysis with these new three groups was performed, only the third group, formed by clusters 2 and 3, showed enrichment in the same categories as in the previous GO analysis, including as a new GO term GO:0009737, response to abscisic acid. Focusing on the genes contributing to that GO term, we observed that, under water stress conditions, HB clearly affects the ABA response since a differential expression of those genes was observed (Supplemental Figure 8), correlating with the lower ABA levels observed (Figure 4E).

### HB application improves productivity under water-limiting conditions

To study whether drought tolerance conferred by HB treatments in tomato plants could contribute to improve productivity, we carried out treatments in an experimental field under limited water availability (50 %).

On the one hand, we examined tomato fruit set throughout the experimental trial. As shown in Figure 6A, plants treated with HB under water stress developed significantly more flowers than non-treated plants during almost the entire experiment. Related to this, we detected a set of genes involved in flower development that were altered by HB treatments under drought conditions in the RNA-seq analysis, which could explain the observed differences (Supplemental Figure 9). In addition, the greater number of flowers translated into a higher fruit set yield at 12 DA-F (Days After treatment F, See Materials and Methods) (Figure 6B). On the other hand, we assessed the number of fruits and total productivity at different time points. At the beginning of the harvest period, the number of fruits on HB-treated plants was lower than on non-treated plants. However, from the fifth harvest (14 DA-H), the number of fruits on HB-treated plants became similar to that of untreated tomato plants. At the end of the harvesting period, HB treated plants produced a higher yield (23%) than non-treated plants (Supplemental Figure 10A). Furthermore, the same trend was observed regarding fruit weight (Supplemental Figure 10B), thus indicating that periodic HB treatments can improve tomato fruit size and productivity under drought conditions. The harvested fruits were also classified depending on their size into extra, class I, class II or class III. Both fruit weight and fruit number were higher in the extra and class I groups, in tomato plants treated with HB subjected to drought conditions (Supplemental Figure 10C and 10D). Our results indicate that HB treatments increase productivity and quality of the fruits in water-stressed tomato plants.

**Figure 6.**
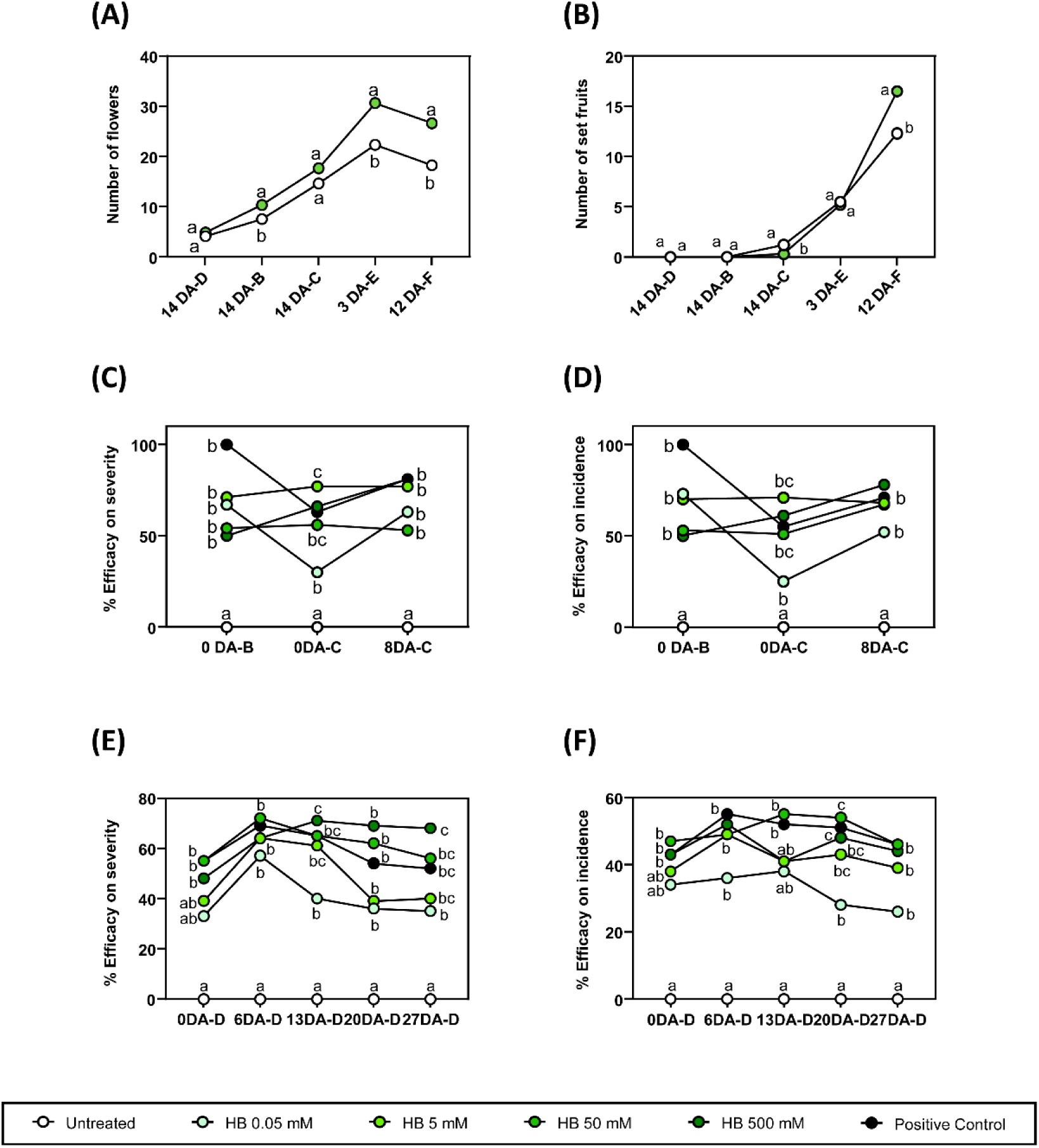
Agricultural applications of HB treatments under open field experiments. Number of flowers (A) and number of set fruits (B) of tomato plants under limited water availability conditions (50%) and treated (+HB) or not (-HB) with 5 mM HB. Time units are referred as follows: X DA Z, being X day (0-18); DA Days After; Z day of treatment (A-H). Points represent mean values. Different letters indicate significant differences at each time point (P < 0.05, two-way ANOVA with Tukey HSD). Efficacy on severity (C) and efficacy on incidence (D) of tomato plants naturally infected by Pseudomonas syringae and either untreated, HB weekly-treated at different concentrations (0.05, 5, 50, 500 mM) or treated with the positive control OXICOOP (0.5%). Time units are referred as follows: X DA Z, being X day (0-8); DA Days After; Z day of treatment (A-C). An ANOVA test was performed, and different letters indicate statistical significances with a P-value < 0.05. Efficacy on severity, (E) and efficacy on incidence (F) of potato plants naturally infected by Phytophthora infestans and either untreated, HB weekly-treated at different concentrations (0.05, 5, 50, 500 mM) or treated with the positive control OROCOBRE (0.3%). Time units are referred as follows: X DA Z, being X day (0-27); DA Days After; Z day of treatment (A-D). An ANOVA test was performed, and different letters indicate statistical significances with a *p* < 0.05.

### HB treatments lead to pathogen resistance

Since HB application activates defense-related genes and confers resistance to the bacteria *Pst* DC3000 under greenhouse conditions with controlled inoculations ^22^, experimental trials were carried out to assess whether HB defensive effects have potential uses in agriculture.

The first trial was performed to confirm the agricultural applications of HB in tomato plants against *Pst* DC3000. For this purpose, tomato plants were treated periodically with different HB concentrations (0.05, 5,10 and 50 mM), using the commercial pesticide OXICOOP 50 (0,5 %) as positive control. The applications started as preventive and were performed as a broadcast foliar application on the tomato plants before appearance of bacterial disease symptoms (see Materials and Methods). On each assessment date, the percentage of leaves presenting symptoms per plot (incidence) and the leaf area infected (severity) were evaluated on 10 plants per plot, and the efficacy of incidence and severity were calculated (Figures 6C and 6D). From the beginning of the experiment, HB-treated plants showed differences with respect to untreated plants in the percentage of efficacy of severity or incidence of the bacterial infection. At 0 DA-C and 8 DA-C intermediate and high concentrations of HB acted in a similar way to the commercial pesticide OXICOOP 50, the efficacy of incidence being between 68 and 78% for high dose of HB treatments and between 77-81% at last assessment considering the percentage of infected leaf.

Furthermore, we tested the efficacy of HB in a different plant-pathogen interaction in which stomatal immunity is crucial, such as potato defense against *Phytophthora infestans*. For this end, the same approach was followed as in the tomato field trial against *Pst* but employing OROCOBRE (0,3%) as a commercial positive control treatment. Regarding the percentage of damaged leaf area, all plots showed statistical differences compared to the untreated control since 6 DA-D. The best results were obtained with the 500 mM HB treatment, reaching efficacies close to 70% and exceeding the performance of the reference OROCOBRE whose efficacy was around 52% at the end of the trial (Figures 6E). Regarding the symptoms in potato plants, HB treatments showed statistical significances from 6 DA-D, obtaining similar efficacy on the incidence of downy mildew than the treatments with the commercial fungicide (Figures 6F).

Therefore, our results indicated that HB can be efficiently used to prevent diseases in agriculture caused by pathogens whose entry is through stomata.

## Discussion

In previous studies, we identified different GLVs that were emitted when tomato plants efficiently resisted bacterial infection with *Pst*. Particularly, the volatile HB has been patented as a universal stomatal closure compound ^30^, but the detailed mechanisms of HB action in stomatal immunity remain undeciphered ^22^. In this work, a transcriptomic analysis revealed that HB treatments triggered the activation of different processes that were related to plant defense mechanisms such as chitin catabolism, glucosamine and aminoglycan metabolism or endopeptidase activity, re-inforcing the role of HB in the defensive response activation (Figure 1A, and Supplemental Figure 1).

The role of stomata as active defensive barriers in plant immunity has been extensively studied. Our results showed that HB treatments induce stomatal closure as effectively as ABA and flg22, (Supplemental Figure 2). Nonetheless, HB treatments in ABA-deficient mutants triggered stomatal closure (Figure 2A), demonstrating that HB-mediated stomatal closure is ABA-independent. Moreover, chemical approaches also indicated that, similarly to flg22, HB treatments induced stomatal closure and plant defense signaling through NADPH-oxidase ROS production and Ca^2+^ ion permeable channels (Figures 2, B, C and D). Furthermore, we have also observed that MPK3 and MPK6 activation is essential for HB-triggered stomatal immunity (Figure 3). Our results correlate with other studies that have shown that GLV exposure induced the activation of these downstream responses. For instance, exposure of Arabidopsis plants to (*E*)-2-hexenal and (*E*)-2-hexenol involved membrane potential depolarization through an increase in cytosolic Ca^2+^ via ROS-activated calcium channels ^31^. Besides, maize and Arabidopsis plants exposed to (*Z*)-3-hexenol and (*E*)-2-hexenal, respectively, increased the transcript abundance of different genes involved in defense signaling ^32,33^. The fact that GLVs are formed from endogenous plant components, are emitted upon pathogen infection, and that their exogenous application activates defense signaling mechanisms suggests that GLVs, and particularly HB, could act as DAMPs ^34^. DAMPs are danger signals released by plants upon pathogen attack that have a relevant role in amplifying the immune responses.

Drought is considered one of the most important factors limiting crop productivity, and new strategies are needed to cope with this global threat. In response to drought stress conditions, the first option for plants is to partially close stomata to prevent water loss through transpiration ^35^. In this study, we have shown that the stomatal aperture ratio of HB-treated tomato plants was similar to the stomatal aperture ratio of plants that were subjected to water stress (Figure 4A), a phenomenon that led to minimize weight loss and ion leakage after 6 days of water deprivation (Figures 4 B and C), indicating that HB-mediated stomatal closure was effective in drought and alleviated its effects. Interestingly, levels of both proline and ABA were lower in HB-treated tomato plants subjected to water deprivation (Figure 4D and 4E), whilst the expression of a *LEA* gene, which is thought to participate in membrane protein stability, osmotic adjustment and macromolecular stabilization ^36^, is highly induced (Supplemental Figure 6C). The transcriptomic analysis of HB-treated plants under drought conditions (Figure 5) appear to indicate that HB-mediated stomata closure could be producing a higher induction of the abiotic responses and a limitation of the water transport, therefore alleviating the drought effect in tomato plants. Besides, the downregulation of the hormone response (Figure 5B) could be associated with the observed ABA repression (Figure 4E), thus reinforcing the idea that HB acts in a ABA independent manner (Figure 2A). In fact, when we compared both RNAseq analyses and we focused on clusters 2 and 3, which only respond to HB in water deficit conditions, an enrichment on the GO corresponding to ABA response was observed, correlating the repression of ABA related genes (Supplemental Figure 8) with the lower levels of ABA observed in those plants (Figure 4E). Our results suggest that HB is conferring tolerance to drought in tomato plants through ABA-independent mechanisms, which appear to be not necessary but replaced by the HB-mediated response.

The application of osmoprotectans and biostimulants as elicitors to confer plant tolerance to different biotic and abiotic stresses through priming is a promising alternative to conventional techniques. The priming mechanism enables plants to respond more robustly and rapidly, which results in an enhanced resistance or tolerance to a stress condition^37^. Most-know priming compounds are synthetic SA analogs, such as 2,1,3-Benzothiadiazole (BTH) or 2,6-dichloroisonicotinic acid and its methyl ester (both referred to as INA) which trigger the activation of systemic acquired resistance (SAR), and the non-protein amino acid β-aminobutyric acid (BABA) also protects plants against various biotic and abiotic stresses in a plethora of plant species ^38^. Many VOCs have also been described as elicitors and priming compounds. Hexanoic acid can activate broad-spectrum defenses by inducing callose deposition and activating SA and JA signaling pathways. Besides, its efficiency has been demonstrated in a wide range of host plants and pathogens ^39,40^. Another example of VOCs as priming agents against abiotic factors is the GLV (*Z*)-3-hexenyl acetate, since maize seeds primed with this volatile exhibited cold tolerance ^41^.

In greenhouse conditions, our laboratory demonstrated that HB produces the induction of resistance against bacterial infection through stomata closure ^22^. In this study, the enhanced resistance to pathogens and tolerance to drought conditions caused by the HB has also been demonstrated in field conditions. Regular exogenous HB treatments in tomato plants under limited water availability conditions resulted in increased number of flowers, which lead to enhanced number and size of tomato fruits compared to non-treated plants (Figures 6A; Supplemental Figure 10). These results are in agreement with the RNA-seq analyses which showed alterations in flowering-related genes in HB-treated tomato plants under drought stress (Figure Supplemental Figure 9). When conditions are unfavorable, plants tend to alter flowering processes to ensure that seed is produced for the next generation ^42^.The observed higher induction of the abiotic responses in HB-treated water-stressed plants (Figure 5) could be affecting the flowering development, therefore explaining the greater number of flowers observed (Figure 6A). An enrichment of GO terms in flowering was not observed in HB-treated plants under drought stress, probably due to the sampling upon HB treatment, containing mostly leaf material and little meristems. Nevertheless, alterations in the expression of flowering-related genes have been specifically detected (Figure Supplemental Figure 9) indicating that HB may influence flower production. Further studies will help us better understand and exploit the effect of HB on flowering and fruit set upon water stress conditions.

HB efficacy in field was also demonstrated in the previously analyzed tomato-bacteria interaction and also against potato late blight caused by *Phytophthora infestans*, the efficacies obtained in both cases being higher than those of commercial agrochemicals (Figures 6C-F) with the added advantage of being natural and non-toxic unlike most conventional pesticides. According to these results, HB emerges as a promising natural elicitor compound against biotic and abiotic stress.

In summary, the results obtained in this work provide evidence for the role of HB in stomatal immunity in tomato and Arabidopsis plants, being ROS generation by NADPH oxidases, Ca^2+^ fluxes, MPK3/6 activation, but not ABA, essential for HB-mediated stomatal immunity. Furthermore, HB-induced stomatal closure resulted in enhanced resistance and tolerance to pathogen infection and drought in field conditions, therefore proposing HB as a novel natural elicitor candidate and a promising strategy for sustainable agriculture.

## Materials and Methods

In this study, tomato (*Solanum lycopersicum*) cv. MoneyMaker was used. Tomato ABA-deficient mutants *flacca (flc)* ^43^ and their corresponding parental Lukullus plants, were kindly provided by Dr. Jorge Lozano (IBMCP, Valencia, Spain). Tomato plants were grown as described in ^22^ For greenhouse drought experiments, water stress was simulated by quitting irrigation during 6 days in the case of stressed plants, whilst control plants were normally watered. HB-treatments were performed every 2 days, until the end of the experiments.

Arabidopsis plants were in Columbia (Col-0) ecotype background. *mpk3* and *mpk6* mutants were kindly provided by Dr. Borja Belda-Palazón (IBMCP, Valencia, Spain). Plants were grown in a chamber under short-day conditions (10/14 h; 19/23 ^◦^C light/darkness), with a relative humidity ranging from 50 to 60%.

### HB treatments under greenhouse conditions

Treatments were carried out on 4-week-old tomato plants either in closed chambers or by spray. Tomato plants were placed into 121 L transparent methacrylate chambers containing hydrophilic cotton swabs soaked with 5 μM of (*Z*)-3-hexenyl butyrate (HB; Sigma-Aldrich, Saint Louis, MO, USA) or distilled water in the case of control plants. Methacrylate chambers were hermetically sealed during the 24 h treatment. In the case of spray treatments, tomato plants were pre-treated by spray with HB at a concentration of 2 mM or distilled water (mock), containing 0.05% Tween 20 as wetting agent.

### Bacterial inoculation assays

The bacterial strain used in this study was *Pseudomonas syringae* pv. tomato (*Pst*) DC3000 (kindly provided by Dr. Selena Gimenez, Centro Nacional de Biotecnología, Madrid, Spain). Bacterial inoculations were performed as described in López-Gresa *et al.* (2018). For spray inoculation assays on Arabidopsis plants, a *Pst* DC3000 culture (OD_600_ = 0.2) was resuspended in 10 mM MgCl_2_ + 0.05% Silwet L-77, and sprayed onto 4-week old plants. The bacterial growth experiments were carried out by sampling three leaf discs (1 cm^2^ each) from each plant at 72 hours post inoculation. Tissue samples were ground in 10 mM MgCl_2_, serial dilutions were done and plated on Petri dishes containing King’s B medium supplemented with rifampicin. Bacterial colonies were counted 48 h after serial dilutions.

### Stomatal aperture measurement

To measure stomatal aperture in tomato plants, clear nail polish was applied in the abaxial part of five leaves from three independent plants. Once the film was dry, it was carefully peeled off and leaf impressions were obtained.

In the case of leaf discs, they were taken from 3- to 4-week old plants and floated on Murashige & Skoog medium (MS) for 3 h under light, in order to induce stomatal opening. Then, stomata closing elicitors (1 μM flg22, 10 μM ABA and 50 μM HB) were added to the medium for 3 h. For chemical inhibitors, 2 mM salicylhydroxamic acid (SHAM), 20 μM diphenyliodonium chloride (DPI), 2 mM ethyleneglycol-bis(β-aminoethyl)-N,N,Nʹ,Nʹ-tetraacetic acid (EGTA) and 20 μM 2-(2-Amino-3-methoxyphenyl)-4H-1-benzopyran-4-one (PD98059) were added 30 minutes before incubation with the elicitors.

Samples were then visualized under a Leica DC5000 microscope (Leica Microsystems GmbH, Wetzlar, Germany) and pictures of different regions were analyzed with *ImageJ* software. Aperture ratio was measured as stomata width/length from at least 50 stomata per plant and/or treatment, considering a value of 1 as a totally opened stoma.

### Ion leakage estimation

Five leaf discs (1 cm^2^) were excised using a stainless steel cork borer, then immersed in tubes containing 40 mL of deionized water and shaken at 200 rpm for 2 h at 28°C. The conductivity of the solution (L1) was measured with a conductivity meter (DDS-11A, Shanghai Leici Instrument Inc., Shanghai, China). After that, the solution containing the leaf discs was boiled for 15 minutes, cooled to room temperature, and the conductivity of the disrupted tissues (L2) was measured. Ion leakage was calculated as the ratio of L1 to L2.

### RNA isolation and RT-qPCR analysis

Total RNA of tomato leaves was extracted as described in López-Gresa *et al.* (2018). Quantitative PCR was carried out as previously described ^44^. The housekeeping gene transcript actin was used as the endogenous reference. The PCR primers were designed using the online service Primer3 (https://primer3.ut.ee) and are listed in Supplemental Table 1.

### RNA-Seq and GO analyses

Six individual MoneyMaker tomato plants of 4 week-old were placed into methacrylate chambers and treated with HB or distilled water as described above. Twenty-four hours post-treatment, leaf samples were collected and total RNA was extracted and analyzed using an 2100 Agilent Bioanalyzer (Agilent Technologies, Inc., Santa Clara, CA, USA) to check RNA integrity and quality. RNA-Seq and bioinformatics analysis were performed by Genomics4All (Madrid, Spain). Samples were sequenced with x50 genome coverage using 1 x 50 bp reads. Raw data was aligned to *Solanum lycopersicum* genome using HISAT2 v2.1.0 ^45^. StringTie was used to obtain the differential expression ^46^. Data and transcript-level expression analysis, and functional enrichment analysis were performed following previously described protocols ^47,48^. GO enrichment analysis were performed with Panther Classification System (https://pantherdb.org) ^49^.

### MPK phosphorylation assay

Leaf discs were excised from 3- to 4-week-old tomato and *Arabidopsis thaliana* plants and floated on MS medium for 3 h under light as described above. Then, 1 μM flg22, 10 μM ABA or 50 μM HB treatments were performed, and samples were taken 15, 30 and 60 minutes after treatments, and immediately frozen in liquid nitrogen. Frozen leaf samples were homogenized and proteins extracts were obtained by Laemmli extraction method. Briefly, leaf disc powder (100 mg) was mixed with 150 µL 2x Laemmli buffer and incubated on ice for 20 minutes, vortexing every 5 minutes. Samples were thereafter boiled for 10 minutes, cooled on ice for 5 minutes, and centrifuged at 12000 x *g* for 5 minutes at 4 °C to clarify homogenates. The resulting supernatants (10 μL) were analyzed by Western Blot with Phospho-p44/42 MAPK (Erk1/2) (Thr202/Tyr204) (Cell Signaling, 1679101S; dilution 1:1,000) primary antibodies, and peroxidase-conjugated goat anti-rabbit IgG (Jacksons, 111035144; dilution 1:20,000) secondary antibodies.

### Open-field HB treatments

#### Abiotic stress

This assay was carried out at Picanya, located in Valencia, Spain (39°26′08″N, 0°26′09″O). Due to the volatility of the compound, the field trial was performed in two separate greenhouses, one for HB treatments and the other one for control treatments. The experimental design used was randomized blocks (9 m^2^) with 4 biological replicates per treatment. Plants were sprayed with 5 mM HB every two weeks. To provoke water stress, irrigation was completely stopped. The soil capacity was continuously monitored and once it reached 50%, the water regime was re-established. Different parameters like number of flowers, fruit set, and total fruits as well as fruit weight were assessed. Moreover, total yield was analyzed at harvest per categories as follows: small size (25-50 mm diameter, 25-100 g weight); medium size (50-75 mm diameter,100-200 g weight); big size (75-100 mm diameter, 200-300 g weight); and extra size (>100 mm diameter, >300 g weight).

In all the field trials, time units are referred as follows: X DA Z, being X day (0-18); DA Days After; Z day of treatment (A-H).

Abiotic stress trials were performed under General Standards, in accordance with EPPO Guidelines PP 1/153(4), PP 1/181(4), PP 1/135 (5) and PP 1/239(2).

#### Biotic stress

The trial of tomato plants inoculated with *Pseudomonas syringae* was performed under open field conditions in Torrellano (Alicante, Spain, 38°17′39″N, 0°35′11″O). The tomato plants were transplanted on August 26^th^, 2020 and the commercial variety Muchamiel was used. The experiment comprised 6 plots (10 plants each) treated with HB at 0.05, 5, 50, and 500 mM, plus another plot that was treated with the commercial pesticide OXICOOP 50 (0,5%) as a positive control, and one plot was left untreated. Treatment applications were carried out using a motorized knapsack sprayer. Three applications were carried out, the application A being preventive and subsequent applications (B and C) were performed within 6 to 8 days intervals coinciding with the persistence of the effect of HB ^22^.

To determine the efficacy of HB for the control of *Downy mildew* in potatoes, an independent open field condition trial was performed in Borbotó (Valencia, Spain, 39°30′55″N, 0°23′27″O). The potato plants were transplanted on February 18^th^ 2019 and the variety used was Vivaldi. The experimental design used was randomized block with 6 plots (one plot per treatment) and 60 plants in each one. Foliar treatments were performed following the same methodology and strategy than in the case of the tomato experiment, but in this case ORO-COBRE ZINEB (0.3%) was used as positive control. Four applications were carried out, being application A before disease appears, and following applications (B, C and D) with 7±1 day intervals.

In both experiments, the degree of infection was assessed as percentage of leaves presenting symptoms (incidence) and percentage of infected leaf area per diseased leaf (severity). The efficacy (E) of the treatments was calculated according to the Abbott’s formula ^50^, using the percentage of leaf area affected in control (C) and treated (T) groups. The following formula was used: E=[(C-T)/C] *100 Tomato and potato-pathogen inoculation trials were performed in accordance with EPPO (Efficacy evaluation of Plant protection Products) guidelines PP1/135(4), PP1/152(4) and PP1/181(4), and specific EPPO PP1/2(4).

## Statistical analysis

The statistical analysis of two or more variables was carried out by using Student’s *t*-test or analysis of variance (ANOVA) respectively, employing the GraphPad’s Prism 9 software. In all the analyses, a *p*-value < 0.05 was considered statistically significant.

## Acknowledgements

We would like to thank Químicas Meristem (Valencia, Spain), especially to Dr. Giovanni Pensabene for his excellent technical support in the open field assays.

## Contributions

MPL-G and PL designed the research; CP, BB-P, FV and JPP performed research; LJ, FV and JPP contributed RNA-Seq analytic tools; CP, BB-P, FV JPP, LJ, IR, JMB, MPL-G and PL analyzed and discussed data; CP, MPL-G and PL wrote the paper and incorporated the input of the rest of the authors.

## Data availability statement

Data available on request.

---

**Supplemental Figure 1. Validation of RNA-seq data by RT-qPCR.** 4 weeks-old tomato plants were treated (+HB) or not (-HB) with 5 μM HB into methacrylate chambers. Gene expression analysis of the *PR1* (X68738), *PR5* (X70787), a pathogen recognition receptor (Solyc12g036793), plant cadmium resistance 2 (Solyc06g066590) and chlorophyll binding genes (Solyc06g069730; Solyc06g069000; Solyc12g011280) was examined by RT-qPCRs using total RNA from leaves. The RT-qPCR values were normalized with the level of expression of the actin gene. Data represent the mean ± SEM of a representative experiment (n=4). Double (**) and triple (***) asterisks indicate significant differences between treatments with *p*-value < 0.01 and *p*-value < 0.001, respectively (Student’s *t*-test).

**Supplemental Figure 2. Dose-response (A) and time course (B) analysis of HB, flg22 and ABA in tomato leaves stomatal aperture.** Data represent the mean ± SEM of a representative experiment (n=50). Different letters indicate statistically significant differences for each treatment at each time point (*p* < 0.05, one-way ANOVA with Tukey HSD Test).

**Supplemental Figure 3. MPK3/6-mediated downstream signaling upon HB treatments.** Real-time qPCR analysis of *WRKY33A* **(A),** *WRKY33B* **(B),** *ETR4* **(C)** and an ethylene responsive factor (*ERF)* **(D)** gene expression in HB-treated plants. Expression levels are relative to tomato mock plants and normalized to the tomato actin gene. Data represent the mean of three independent plants ± SEM. Asterisk (*) and double asterisks (**) indicate statistically significant differences with *p* ≤ 0.05 and *p* ≤ 0.01, respectively (Student’s *t*-test). **(E)** Heat map of *WRKY33A*, *WRKY33B*, *ETR4* and an *ERF* from the RNA-seq from watered plants experiment. The numbers in each box represent the average of the log_2_ of the fragments per kilobase of exon per million mapped fragments (FPKM) of each sample.

**Supplemental Figure 4. Effect of HB treatments on chlorophyll content in tomato leaves.** Chlorophyll a, chlorophyll b and total chlorophyll content in tomato plants treated (+HB) or not (-HB) with HB, in normal (+H_2_O) or water stressed conditions (-H_2_O). Data correspond to six independent plants ± SEM of a representative experiment. No statistically significant differences were found.

**Supplemental Figure 5. Gene expression in tomato plants treated with HB under drought conditions**. Tomato plants were periodically sprayed (+HB) or not (-HB) with 2 mM HB, and plants were subjected (-H_2_0) or not (+H_2_0) to 6 days of water deprivation. Gene relative expression of *P5CS1* (Solyc06g01970) **(A)**, 9-cis-epoxycarotenoid dioxygenase (Solyc07g056570) **(B),** zeaxanthin epoxidase (Solyc02g090890) **(C)**, and ABA 8’-hydroxylase (Solyc04g078900) **(D)** genes. RT-qPCR values were normalized with the level of expression of the actin gene. Bars represent the mean ± SEM of a representative experiment. Different letters indicate statistically significant differences for each genotype and treatment (*p* < 0.05, two-way ANOVA with Tukey HSD).

**Supplemental Figure 6. ABA-related gene expression profiles in tomato plants treated with HB under drought conditions.** Plants were treated with HB (+HB) or water (-HB) periodically. Water-stressed plants were subjected to water deficit for 6 days. *RAB18* **(A),** *MYB44* **(B)** and *LEA* family **(C)** gene relative expression in plants under normal (+H_2_0) or water deficit conditions (-H_2_0). Gene expression analysis were examined by RT-qPCRs using total RNA from leaves. The RT-qPCR values were normalized with the level of expression of the actin gene. Bars represent the mean ± the standard deviation of a representative experiment (n=3). Different letters indicate statistically significant differences for each treatment and condition (*p* < 0.05, one-way ANOVA with Tukey HSD Test).

**Supplemental Figure 7. Cluster analysis of RNAseq data of tomato plants under different treatments (+/-HB) and conditions (+/-H_2_0).** Plants were treated (+HB) or not (-HB) with 2 mM HB and were subjected to water deprivation (-H_2_0) for 6 days or normally watered (+H_2_0). Blue lines correspond to HB treatments relative expression in water conditions, and black lines represent expression data from plants subjected to drought.

**Supplemental Figure 8. Heat map of ABA-related genes from the RNA-seq from non-watered plants experiment.** The numbers in each box represent the average of the log_2_ of the fragments per kilobase of exon per million mapped fragments (FPKM) of each sample.

**Supplemental Figure 9. Heat map of flowering related genes from the RNA-seq from non-watered plants experiment.** The numbers in each box represent the average of the log_2_ of the fragments per kilobase of exon per million mapped fragments (FPKM) of each sample.

**Supplemental Figure 10. HB treatments in tomato plants improve productivity under drought conditions in open field experiments.** Total harvested fruits **(A)** and mean fruit weight **(B)** of tomato plants under limited water availability conditions (50%) and treated (+HB) or not (-HB) with 5mM HB. Time units are referred as follows: X DA Z, being X day (0-18); DA Days After; Z day of treatment (A-H). Points represent mean values. Different letters indicate significant differences at each time point (*p* < 0.05, two-way ANOVA with Tukey HSD). Number of harvested fruits **(C)** and total weight of harvested fruits **(D)** per categories were analyzed as follows: Class III-small size (25-50 mm diameter, 25-100 g weight); Class II-medium size (50-75 mm diameter,100-200 g weight); Class I-big size (75-100 mm diameter, 200-300 g weight); and Extra size (>100 mm diameter, >300 g weight).

**Supplemental Table 1.** Primer sequences used for quantitative RT-PCR analysis of the selected tomato genes.

