## Supplemental Information for "Signaling mechanisms and agricultural applications of (*Z*)-3-Hexenyl Butyrate-mediated stomatal closure"

Supplemental Figure 1

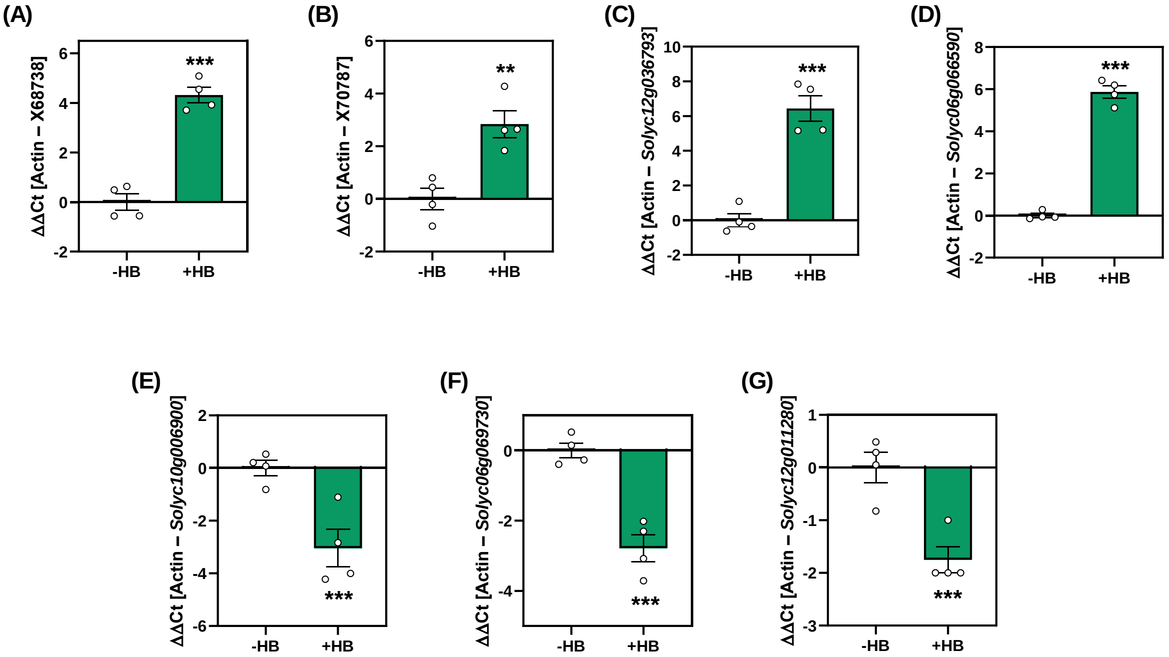


**Supplemental Figure 1**. **Validation of RNA-seq data by RT-qPCR.** 4 weeks-old tomato plants were treated (+HB) or not (-HB) with 5 μM HB into methacrylate chambers. Gene expression analysis of the *PR1* (X68738), *PR5* (X70787), a pathogen recognition receptor (Solyc12g036793), plant cadmium resistance 2 (Solyc06g066590) and chlorophyll binding genes (Solyc06g069730; Solyc06g069000; Solyc12g011280) was examined by RT-qPCRs using total RNA from leaves. The RT-qPCR values were normalized with the level of expression of the actin gene. Data represent the mean ± SEM of a representative experiment (n=4). Double (**) and triple (***) asterisks indicate significant differences between treatments with *p*-value < 0.01 and *p*-value < 0.001, respectively (Student's *t*-test).

### Supplemental Figure 2

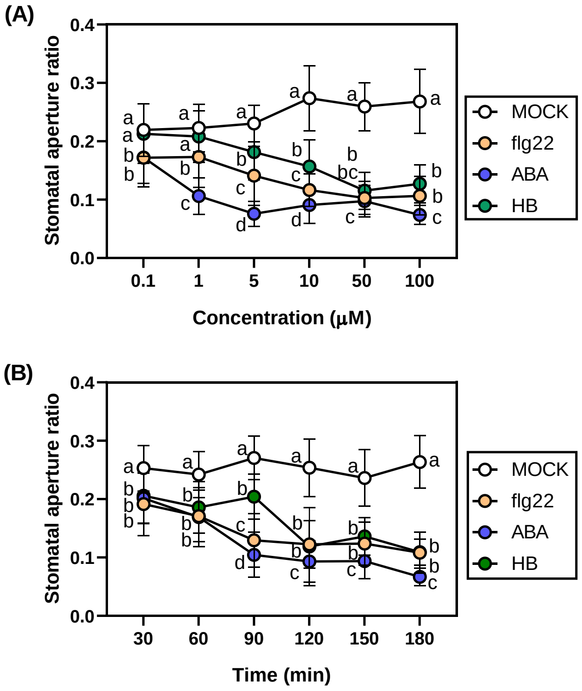


**Supplemental Figure 2. Dose-response (A) and time course (B) analysis of HB, flg22 and ABA in tomato leaves stomatal aperture.** Data represent the mean ± SEM of a representative experiment (n=50**).** Different letters indicate statistically significant differences for each treatment at each time point ($p < 0.05$, one-way ANOVA with Tukey HSD Test).

### Supplemental Figure 3

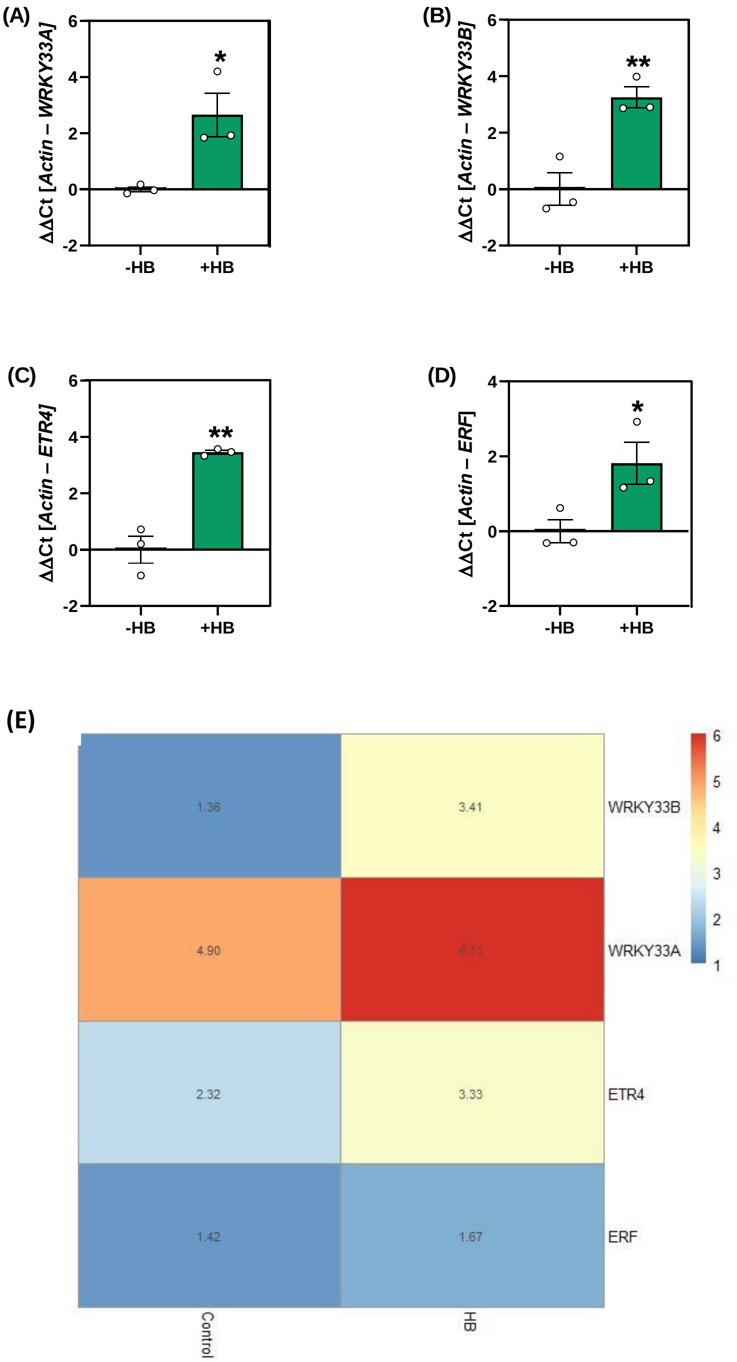


**Supplemental Figure 3. MPK3/6-mediated downstream signaling upon HB treatments.** Real-time qPCR analysis of *WRKY33A* **(A),** *WRKY33B* **(B),** *ETR4* **(C)** and an ethylene responsive factor (*ERF)* **(D)** gene expression in HB-treated plants. Expression levels are relative to tomato mock plants and normalized to the tomato actin gene. Data represent the mean of three independent plants ± SEM. Asterisk (*) and double asterisks (**) indicate statistically significant differences with $p \leq 0.05$ and $p \leq 0.01$, respectively (Student's *t*-test). **(E)** Heat map of *WRKY33A*, *WRKY33B*, *ETR4* and an *ERF* from the RNA-seq from watered plants experiment. The numbers in each box represent the average of the $\log_2$ of the fragments per kilobase of exon per million mapped fragments (FPKM) of each sample.

Supplemental Figure 4

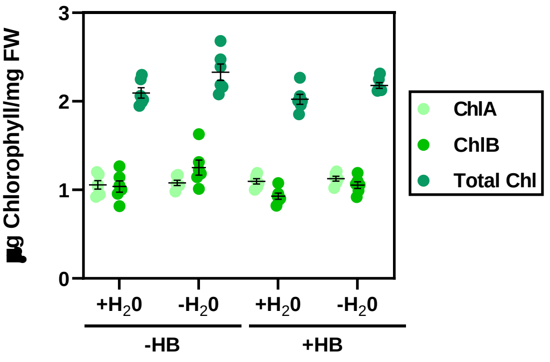


**Supplemental Figure 4. Effect of HB treatments on chlorophyll content in tomato leaves.** Chlorophyll a, chlorophyll b and total chlorophyll content in tomato plants treated (+HB) or not (-HB) with HB, in normal (+$H_2O$) or water stressed conditions (-$H_2O$). Data correspond to six independent plants ± SEM of a representative experiment. No statistically significant differences were found.

Supplemental Figure 5

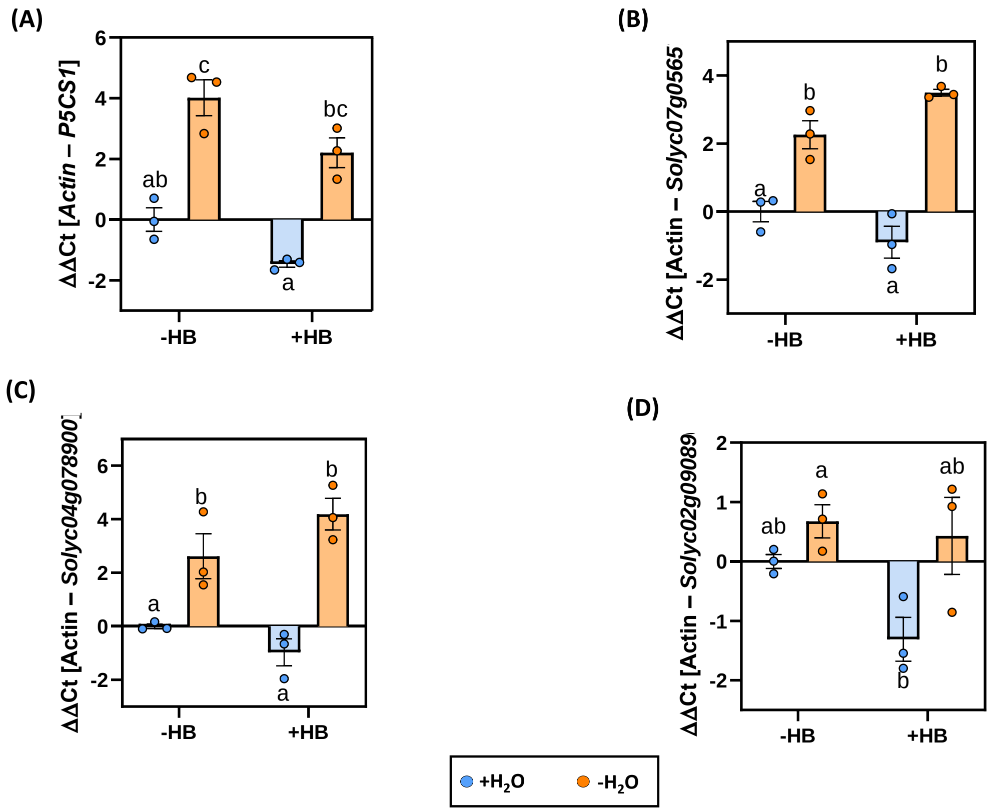


**Supplemental Figure 5. Gene expression in tomato plants treated with HB under drought conditions**. Tomato plants were periodically sprayed (+HB) or not (-HB) with 2 mM HB, and plants were subjected (-$H_20$) or not (+$H_20$) to 6 days of water deprivation. Gene relative expression of *P5CS1* (Solyc06g01970) **(A)**, 9-cis-epoxycarotenoid dioxygenase (Solyc07g056570) **(B)**, zeaxanthin epoxidase (Solyc02g090890) **(C)**, and ABA 8'-hydroxylase (Solyc04g078900) **(D)** genes. RT-qPCR values were normalized with the level of expression of the actin gene. Bars represent the mean ± SEM of a representative experiment. Different letters indicate statistically significant differences for each genotype and treatment ($p < 0.05$, two-way ANOVA with Tukey HSD).

Supplemental Figure 6

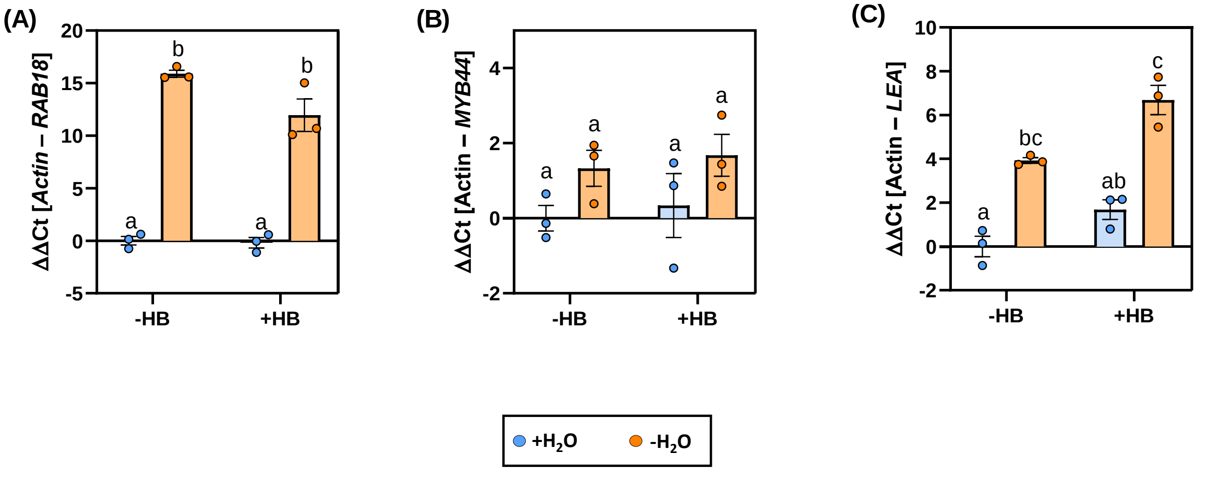


**Supplemental Figure 6. ABA-related gene expression profiles in tomato plants treated with HB under drought conditions.** Plants were treated with HB (+HB) or water (-HB) periodically. Water-stressed plants were subjected to water deficit for 6 days. *RAB18* **(A),** *MYB44* **(B)** and *LEA* family **(C)** gene relative expression in plants under normal (+$H_2$0) or water deficit conditions (-$H_2$0). Gene expression analysis were examined by RT-qPCRs using total RNA from leaves. The RT-qPCR values were normalized with the level of expression of the actin gene. Bars represent the mean ± the standard deviation of a representative experiment (n=3). Different letters indicate statistically significant differences for each treatment and condition ($p < 0.05$, one-way ANOVA with Tukey HSD Test).

Supplemental Figure 7

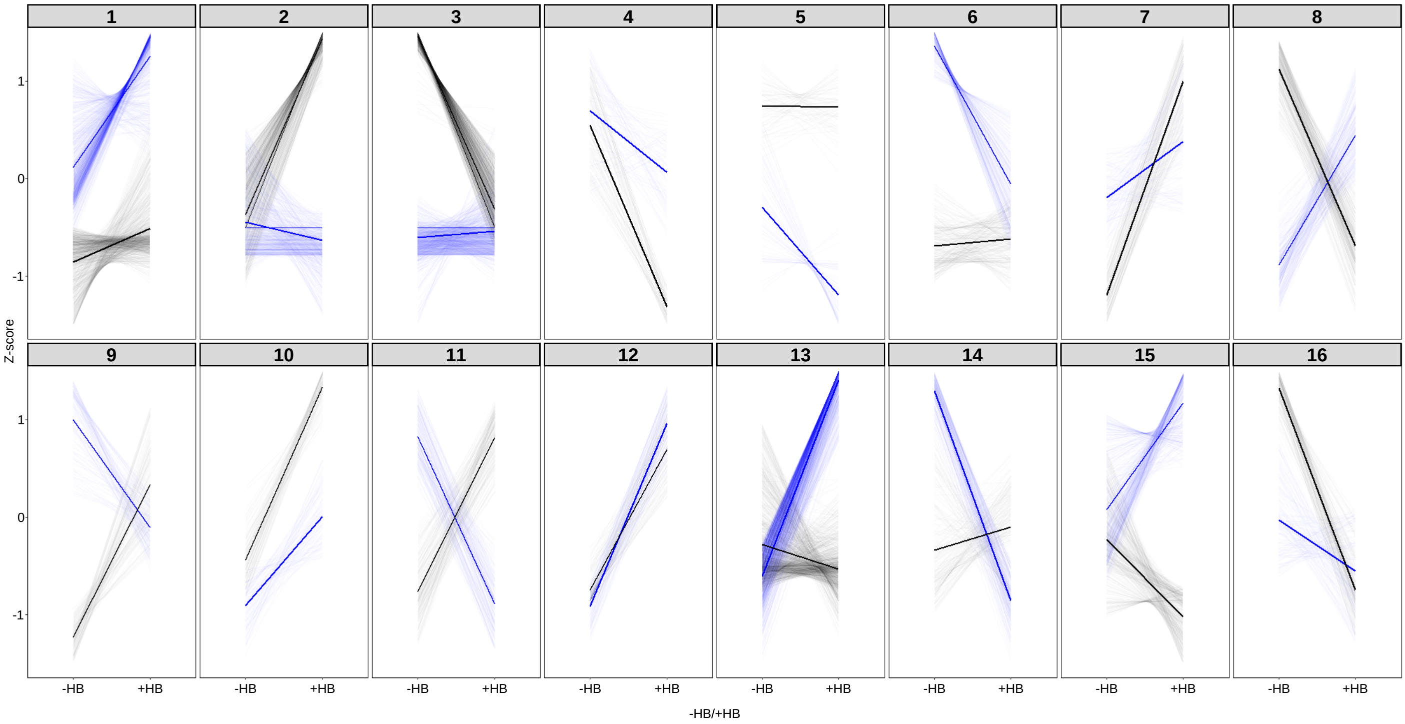


**Supplemental Figure 7. Cluster analysis of RNAseq data of tomato plants under different treatments (+/-HB) and conditions (+/-$H_20$).** Plants were treated (+HB) or not (-HB) with 2 mM HB and were subjected to water deprivation (-$H_20$) for 6 days or normally watered (+$H_20$). Blue lines correspond to HB treatments relative expression in water conditions, and black lines represent expression data from plants subjected to drought.

Supplemental Figure 8

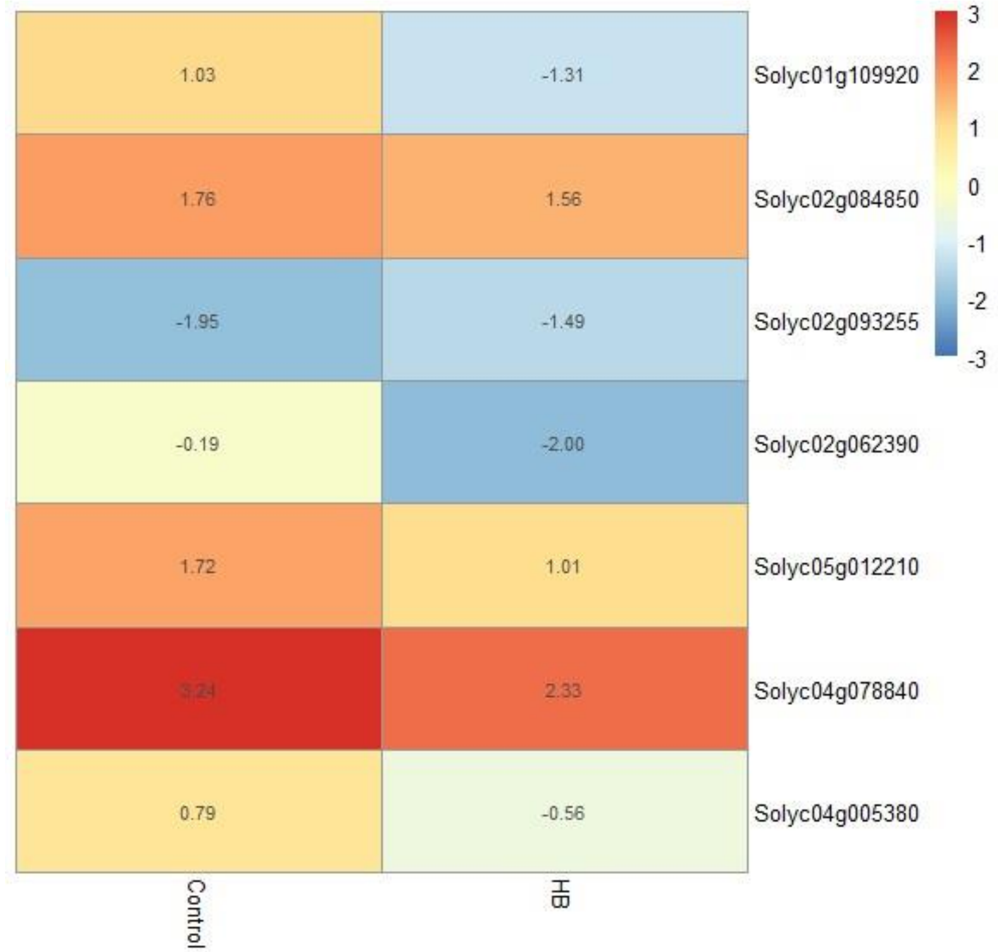


**Supplemental Figure 8. Heat map of ABA-related genes from the RNA-seq from non-watered plants experiment.** The numbers in each box represent the average of the $log_2$ of the fragments per kilobase of exon per million mapped fragments (FPKM) of each sample.

Supplemental Figure 9

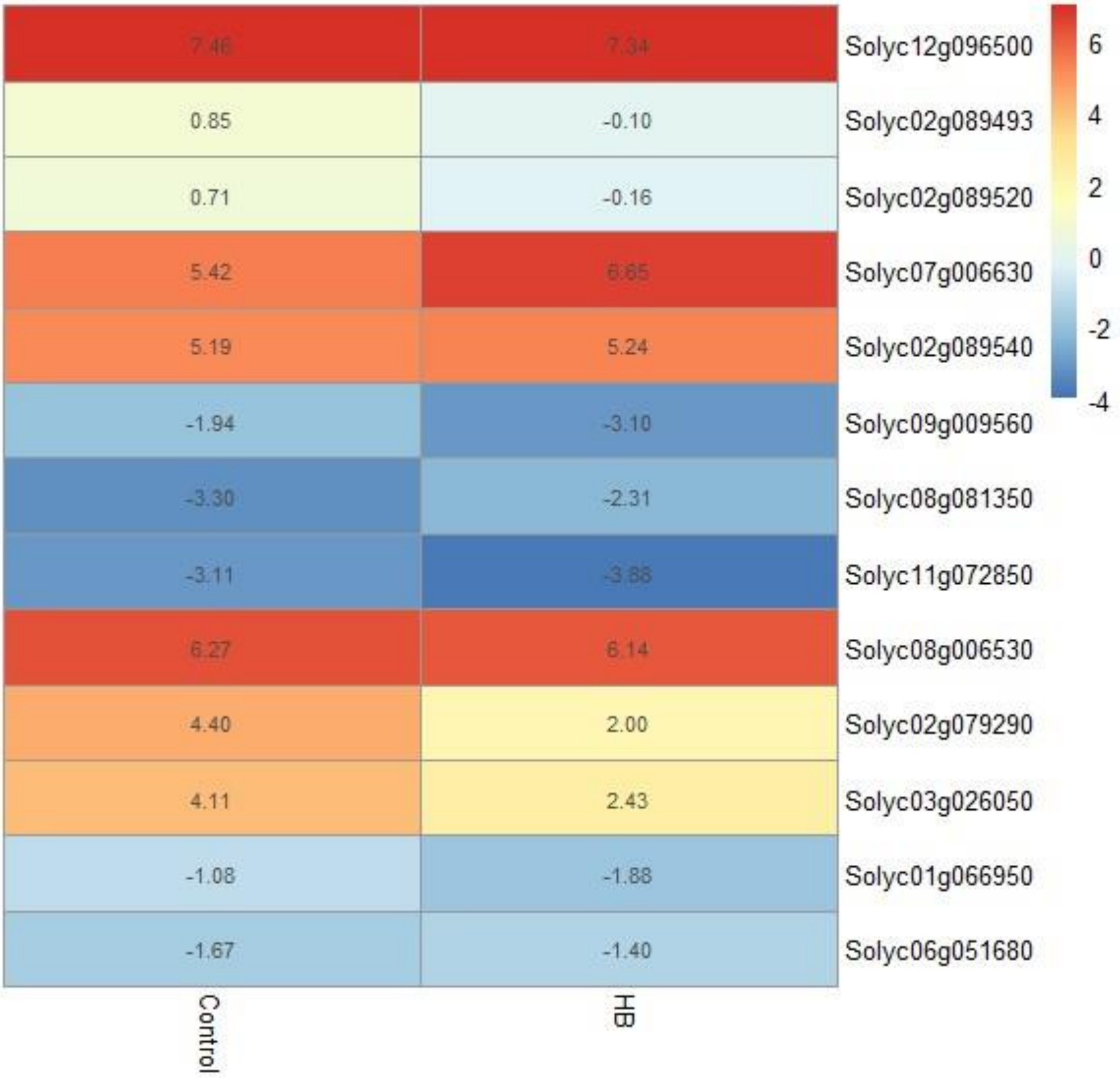


**Supplemental Figure 9. Heat map of flowering related genes from the RNA-seq from non-watered plants experiment.** The numbers in each box represent the average of the $log_2$ of the fragments per kilobase of exon per million mapped fragments (FPKM) of each sample.

Supplemental Figure 10

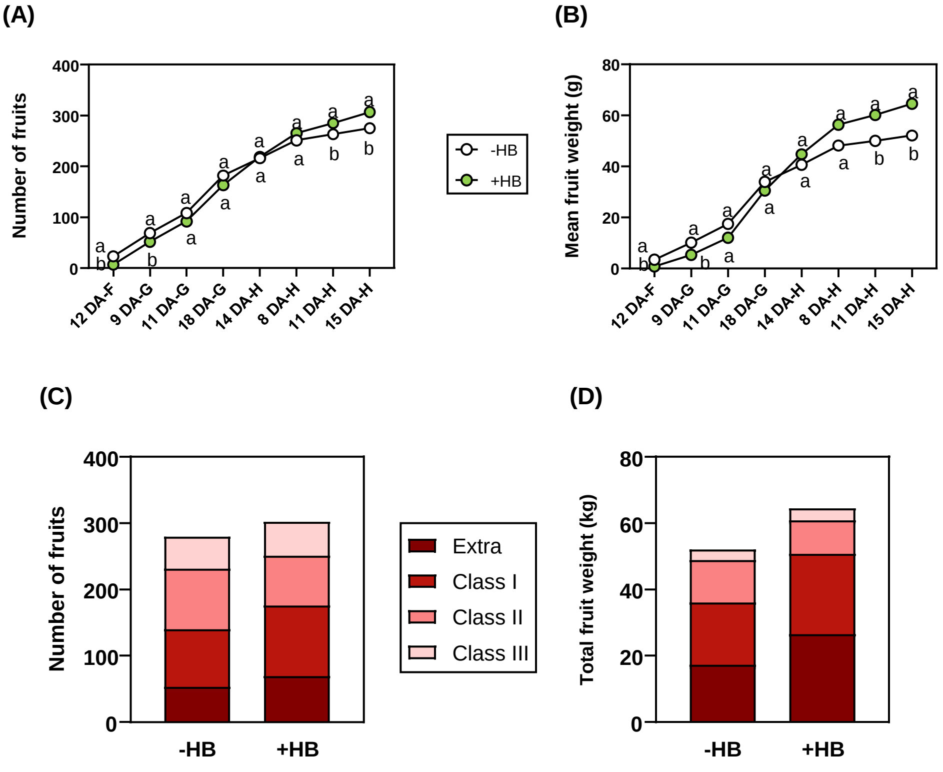


**Supplemental Figure 10. HB treatments in tomato plants improve productivity under drought conditions in open field experiments.** Total harvested fruits **(A)** and mean fruit weight **(B)** of tomato plants under limited water availability conditions (50%) and treated (+HB) or not (-HB) with 5mM HB. Time units are referred as follows: X DA Z, being X day (0-18); DA Days After; Z day of treatment (A-H). Points represent mean values. Different letters indicate significant differences at each time point ($p$ < 0.05, two-way ANOVA with Tukey HSD). Number of harvested fruits **(C)** and total weight of harvested fruits **(D)** per categories were analyzed as follows: Class III-small size (25-50 mm diameter, 25-100 g weight); Class II-medium size (50-75 mm diameter,100-200 g weight); Class I-big size (75-100 mm diameter, 200-300 g weight); and Extra size (>100 mm diameter, >300 g weight).

Supplemental Table 1

| **Gene** | **Forward primer (5'-3')** | **Reverse primer (5'-3')** |
|---|---|---|
| *Actin* | CTAGGGTGGGTTCGCAGGAGATGATGC | GTCTTTTTGACCCATACCCACCATCACAC |
| *ERF* | GTTTGAGGCCCAAAAGGAA | TCAAGTGGCGTTCTCAACAG |
| *ETR4* | CTGCAGATTGGAATGAATGG | ATAAGGCACCGTCAACATCA |
| *MYB44* | GGGAGTCGAACAAAAGCAAC | TCCAAGCCCTAAACCACTTG |
| *LEA* | TTCCCTGCTGACAAAGCTAGAGCA | AGGGTGCAGATGAAACTGATCCGA |
| *P5CS1* | ACCTTAATCTGGAGGCTTGA | AATTATTTCACCCACCTGCC |
| *RAB18* | CCTGGGATGCATTGAACACC | CACGGGACACCATAACACAC |
| *Solyc12g036793* | TCATTGCCCTTGTGTTCTTG | TGTTTGGGTCTGCCACATTA |
| *Solyc06g066590* | ATGACGGCGTTCTTACCTTG | GCAGGCATCATCAGCATGTA |
| *Solyc10g006900* | ACCACGAAGAGACTGGCATC | ATGGAGGGAAAAGGAGCCTA |
| *Solyc06g069730* | CCCTGGAAGTGTGAACCAAG | TGCAAAGTTGAGTGGGTTGA |
| *Solyc12g011280* | TAGGCCCATTCTCTGGTGAG | GCAAAAGTTTCAGGGTCAGC |
| Solyc02g090890 | GGTGGGTTAGTGTTTGCTTTG | GGTGGGTTAGTGTTTGCTTTG |
| Solyc07g056570 | TACGTTCGAAACGGAGCTAAC | CGTAACTAGCCGACCCATTT |
| Solyc04g078900 | CTCAAGACCCTAATGCCTTCTT | CATCACACATGGACAACCTAGA |
| X68738 | ACTCAAGTAGTCTGGCGCAACTC | AGTAAGGACGTTGTCCGATCGAGT |
| X70787 | TTCGAGGTACGCAACAACTG | ATGCATTGATGACCCATGTTT |
| *WRKY33A* | GCATTACTGTCAACCATCGC | AACTTCGCGGATTCTCACTT |
| *WRKY33B* | CCACAACAGTCTGAAATGGG | CAGCAAAGCAATGACTCCAT |

**Supplemental Table 1.** Primer sequences used for quantitative RT-PCR analysis of the selected tomato genes.
